# EXOSC3 G191 Variants Trigger System-Wide Recalibration of RNA Processing Machinery

**DOI:** 10.64898/2026.01.26.701775

**Authors:** H.R. Sagara Wijeratne, Avery M. Runnebohm, Monica P. Barron, Ariana J. Hampton, Rudong Li, Sarah A. Peck Justice, Sophia H. Lenceski, James D. Rooney, Whitney R. Smith-Kinnaman, Gina S. Chang, Emma H. Doud, Jungsu Kim, Stephane Pelletier, Yunlong Liu, Jonah Z. Vilseck, Amber L. Mosley

**Affiliations:** Department of Biochemistry, Molecular Biology, & Pharmacology, Indiana University School of Medicine, Indianapolis, IN 46202; Stark Neurosciences Research Institute, Indiana University School of Medicine, Indianapolis, IN 46202; Department of Medical & Molecular Genetics, Indiana University School of Medicine, Indianapolis, IN 46202; Medical Scientist Training Program, Indiana University School of Medicine, Indianapolis, IN 46202; Indiana University Simon Comprehensive Cancer Center, Indiana University School of Medicine, Indianapolis, IN 46202; Center for Computational Biology & Bioinformatics, Indiana University School of Medicine, Indianapolis, IN 46202; Department of Biology, Marian University, Indianapolis, IN 46222

## Abstract

Pathogenic variants in EXOSC3, a noncatalytic subunit of the RNA exosome, cause pontocerebellar hypoplasia type 1B (PCH1B), yet substantial variability in disease severity is observed among individuals carrying different EXOSC3 alleles. The molecular mechanisms of RNA exosome dysfunction in individuals carrying EXOSC3 p.G191 variants remains unresolved. To address this, we generated CRISPR/Cas9-engineered human cell models harboring EXOSC3 p.G191 variants and performed integrated transcriptomic, proteomic, and computational structural analyses. EXOSC3 p.G191 variants produced extensive, allele- and dosage-dependent alterations in gene expression and splicing, with heterozygous variants causing broad but attenuated disruption relative to homozygous EXOSC3 G191D/G191D cells. All EXOSC3 G191 variants promoted increased skipping of exon 3 in EXOSC3 transcripts, generating a short isoform predicted to encode an unstable proteoform. Molecular dynamics and λ-dynamics simulations predicted substantial thermodynamic destabilization of all EXOSC3 G191 variant proteoforms, consistent with reduced protein abundance and thermal stability measured by global proteomics and PISA. At the protein complex level, EXOSC3 p.G191 variants were associated with coordinated decreases in all RNA exosome core subunits and the exonuclease EXOSC10, consistent with destabilization of RNA exosome assembly and orphan protein decay. In contrast, the catalytic exonuclease DIS3 and multiple independent RNA processing pathways were upregulated, indicating compensatory recalibration of RNA metabolism. Together, these findings link variant-induced alternative splicing, RNA exosome complex destabilization, and adaptive network responses to phenotypic variability in EXOSC3 p.G191-associated disease.

## Introduction

Pontocerebellar hypoplasias (PCH) are a group of rare, genetically heterogeneous neurodegenerative disorders, frequently with prenatal onset, that cause severe degeneration of the pons and cerebellum. Hallmark features of severe PCH include profound motor and cognitive impairments, microcephaly, progressive neurological decline, and/or significant developmental delays. Multiple subtypes of PCH have been identified, each associated with mutations in distinct genes and a range of clinical presentations. Interestingly, many PCH-associated genes encode proteins involved in RNA metabolism and protein synthesis. Mutations in TSEN54, TSEN2, TSEN34, and TSEN15, which encode subunits of the tRNA (transfer RNA) splicing endonuclease complex, are associated with PCH types 2, 4, and 5 ^1–8^. TOE1 (PCH7), encodes an RNA deadenylase required for maturation of RNA Polymerase II produced small nuclear RNA (snRNAs); and CLP1 (PCH10), encodes an RNA kinase involved in tRNA splicing and RNA Polymerase II transcribed mRNA 3’ end processing^9–14^. Several aminoacyl-tRNA synthetases have also been implicated, including DARS2, EARS2, MARS2, QARS, SEPSECS, and RARS2. Of particular relevance to our work, variants in genes encoding several subunits of the RNA exosome complex have been linked to PCH, including EXOSC3 (PCH1B), EXOSC8 (PCH1C), EXOSC9 (PCH1D), and provisionally EXOSC1 (PCH1F)^15–25^. Other subunits of the RNA exosome have also been implicated in neurodegenerative / neurodevelopmental disorders including EXOSC5 and EXOSC2^26–30^. RNA exosome activity is essential in all eukaryotes that have been assessed, including humans, where the core RNA exosome subunits EXOSC1-10, the DIS3 exonuclease, and the MTR4 RNA exosome RNA helicase have all been identified as common essential in the Dependency Map (DepMap) project ^31,32^. The RNA exosome serves fundamental roles in diverse processes including RNA quality control, processing for RNA maturation, and regulated RNA turnover (reviewed in PMID: 26726035, 38637632, 37384835, 30848101). Overall, the breadth of genes associated with PCH has revealed a critical role for RNA processing enzymes and cofactors in addition to tRNA synthetases in cerebellar and pontine development^33–35^.

Within the PCH spectrum, PCH1B is a subtype characterized by marked hypotonia, progressive cerebellar atrophy, and motor neuron dysfunction and caused by variants in the *EXOSC3* gene^15,18,21–23,25^. The extent of PCH1b clinical symptoms is largely dependent on the genotype-phenotype correlation that have been observed in individuals with *EXOSC3* mutations. For example, patients harboring the *EXOSC3* p.Gly31Ala (p.G31A) variant present with absent tendon reflexes, significant neurogenic muscle atrophy, and survival typically limited to the first six months of life^25,36^. However, patients with homozygous *EXOSC3* p.Asp132Ala (p.D132A) variant can live up to the second decade of life^37^. These observations underscore the intricate relationship between specific genetic variants and their impact on RNA metabolism pathways critical for neurodevelopment and/or neuronal survival. *EXOSC3* encodes a core structural component of the RNA exosome complex and contains an RNA binding S1 domain, although weak RNA binding is displayed *in vitro* outside of the context of the full complex^38^. The catalytic activities of the RNA exosome are mediated by associated 3’-5’ exonucleases EXOSC10 and DIS3 or accessory factors such as the RNA helicase MTR4, while core components including EXOSC3 provide structural stability and facilitate substrate recognition and/or RNA threading through the protein complex core^39–43^. In addition to RNA turnover, the RNA exosome has specialized functions in RNA splicing and the regulation of AU-rich element (ARE)-containing transcripts^44–47^. The RNA exosome also plays central roles in ribosomal RNA processing (rRNA) and defects in rRNA processing have been identified in yeast and fly models expressing EXOSC3 variants^15,48^. In our recent studies, we have also observed rRNA processing defects in human cells lines expressing EXOSC3 variants although in an allele-specific manner^47^. The RNA exosome complex also ensures precise intron removal and spliceosomal RNA processing, critical steps in the maturation of protein-coding transcripts^45^. Dysregulation of these processes can lead to widespread splicing defects that could contribute to the pathogenesis of disorders like PCH1B. Through its regulation of many types of cellular transcripts, the RNA exosome not only maintains cellular homeostasis but also influences cellular responses to stress and developmental signals^49–51^. Disruptions in these RNA processing pathways can shed light on how disease associated sequence variants in *EXOSC3* alter mammalian cell homeostasis when compared to isogenic control cell lines. Additionally, our findings can identify how EXOSC3 variants may contribute to the cellular phenotypes and clinical features observed in PCH1B.

Here, we focus on variants of the *EXOSC3* p.G191 codon that have been identified in patients but which have not been previously studied at the molecular level in mammalian cells. *EXOSC3* p.G191 is located at a key structural position at the end of the S1-domain just prior to a linker region in front of the KH-type RNA-binding domain^52^. These domains contribute to EXOSC3 RNA recognition and, given the p.G191 position, p.G191 could contribute to changes in EXOSC3 three-dimensional structural which could influence the overall RNA exosome complex. Interestingly, PCH1B patients who are carriers of compound heterozygous EXOSC3 D132A/G191D variants present with milder disease, without involving the pons, and show intrafamilial variability^23,53,54^. The basis for the higher phenotypic variability in EXOSC3 p.G191 carriers has not been determined at the molecular level. In this study, we utilized CRISPR/Cas9 to generate HEK293T cell models with G191 variants in *EXOSC3* and performed paired multi-omics experimental analyses with computational *in silico* investigations of protein structural dynamics. We show that the transcriptome of *EXOSC3* p.G191 cell lines shows extensive changes in transcript levels along with potentially compensatory increases in RNA processing related factors including DIS3. Surprisingly, G191 variants trigger an alternative splicing event in EXOSC3 leading to an increased frequency of exon 3 skipping. As a consequence, G191 homozygous and heterozygous variants lead to altered EXOSC3 transcript isoform expression including a short form that we show is fully destabilized through molecular dynamics simulations. Exon 3-containing proteoforms with either G191D or G191C variants are predicted to cause thermodynamic folding changes of several kcal/mol as determined by λ-dynamics simulations. G191 variant protein isoforms (proteoforms) lead to widespread alterations in both the transcriptome and the proteome suggesting that EXOSC3 G191 variants cause RNA exosome functional disruption in both neuronal and non-neuronal cell types. The involvement of an alternative splicing event associated with G191 variants could explain the higher degree of disease heterogeneity observed in *EXOSC3* p.G191 carriers.

## Results

### EXOSC3 G191 variants show extensive transcriptome changes

A critical barrier to understanding PCH1B is decoupling the primary molecular consequences of EXOSC3 variants from the secondary cascades of neurodegeneration. While PCH1B manifests clinically in the nervous system, the RNA exosome is a universally essential complex. Therefore, we utilized HEK293T cells to perform a comprehensive structural and functional analysis in a system uncompromised by the apoptosis that occurs in neurons. This approach allows us to define the fundamental, system-wide perturbations in RNA processing machinery and protein stability that precede and likely drive tissue-specific pathogenesis. Although patient fibroblasts and non-affected control fibroblasts have been analyzed in prior work^55^, we have prioritized characterization of EXOSC3 variants in mammalian cells relative to an isogenic control to also lessen challenges associated with differences in genetic background and allow for comparison of functional differences in EXOSC3 variants^47^. Together, these findings showed the importance of including an isogenic control for full interpretation of RNA exosome functional changes. Consequently, to explore the functional effects of the G191 variants on EXOSC3, we utilized CRISPR/Cas9-mediated genome editing to create single clonal HEK293T cell lines expressing the G191 variants and clonal control lines that were also exposed to transient CRISPR/Cas9 expression. This approach allowed us to introduce specific variants at the EXOSC3 locus, enabling a direct examination of the impact of these variants on cellular function relative to isogenic controls. Genomic modifications for each clone were confirmed through Sanger sequencing, which verified the successful introduction of p.G191C or p.G191D variants in EXOSC3. We successfully generated heterozygous G191C/+, heterozygous G191D/+, and homozygous G191D/G191D clonal cell lines, alongside single-cell isolated parental controls. Ultra-deep transcriptomic analysis of the cell lines showed a triploid allele frequency in the G191C heterozygous cell line with two copies of EXOSC3 G191C and one of the parental EXOSC3 sequence (**Fig. S1A**). Since heterozygous carriers of single copy EXOSC3 variants reported to date are unaffected, the inclusion of the EXOSC3 G191D/+ and G191C/+ heterozygous cell lines facilitated an in-depth molecular characterization of variant dosage for G191 related RNA exosome disruption. For subsequent analyses, four paired cell growths were used for multi-omics experiments to minimize batch effects between omics experiments.

To investigate the functional consequences of the p.G191 variants on transcriptome regulation, we performed total RNA sequencing (RNA-seq) followed by computational and statistical analyses. Specifically, we isolated total RNA from HEK293T cells for each genotype along with isogenic parental controls (hence control), for bulk total RNA-seq. This approach allowed us to evaluate both coding and non-coding RNAs, providing insight into how each variant affects global RNA homeostasis. We analyzed *n* = 4 biological replicates per genotype and sequenced at an average depth of 259.82 ± 38.1 million (mean ± SD) reads per sample. This sequencing depth achieved ultra-deep RNA-Seq coverage sufficient for both RNA expression and differential alternative splicing analysis to aid in transcript isoform detection^56^. Statistical analysis for differentially abundant transcripts was performed using the R package DESeq2, applying a Wald test for each variant relative to the control cell line, followed by Benjamini-Hochberg correction of *p*-values^57^. After filtering out lowly expressed transcripts across all samples, we quantified 45,109 transcripts, including 18,915 protein-coding genes, 14,772 long non-coding RNAs, 6,414 pseudogenes (processed and unprocessed), 831 microRNAs (miRNAs), 744 small nuclear RNAs (snRNAs), and 317 small nucleolar RNAs (snoRNAs).

Our transcriptomic analyses revealed widespread changes in RNA abundance across all *EXOSC3* p.G191 genotypes, with the most pronounced effects observed in homozygous G191D/G191D with 7,880 transcripts being significantly changed using an adjusted p-value cutoff of ≤ 0.05 (**Figure 1A**). EXOSC3 G191D/G191D cells surprisingly showed lower total numbers of increased transcripts (1,780) than decreased transcripts (6,100). EXOSC3 G191D/+ heterozygous cells showed differential abundance of 2,938 transcripts with increased abundance and 3,443 with decreased abundance (**Figure S1B**). EXOSC3 G191C/+ cell lines exhibited increased abundance of 3,229 transcripts and decreased abundance of 3,602 transcripts (**Figure S1C**). Principal component analysis of the average transcript abundance for each genotype revealed strong genotype specific effects with control vs. variant separating in PC1 (46% variance) and G191D vs. G191C genotypes separating in PC2 (24% variance, **Fig. S1D**). Heterozygous EXOSC3 G191D/+ samples were close but distinctly separated from G191D/G191D samples, suggesting that many of the genotype-based transcript alterations were observed in both the homozygous and heterozygous clonal cell lines. We have also performed total RNA-Seq analysis of homozygous EXOSC3 D132A/D132A HEK293T cells relative to controls, which revealed more modest changes to the transcriptome than observed for the EXOSC3 G191 cell lines^47^. Direct comparison of the differential transcripts identified in EXOSC3 G191D/G191D and/or D132A/D132A cells showed a higher degree of unique transcript changes for each genotype relative to the number that overlap, suggesting that the transcriptomes are altered in many allele-specific ways. A total of 455 transcripts were shared between the two datasets and pathway analysis revealed significant enrichment in transcripts of proteins of medium expression levels associated with the cerebellum and its white matter processes (HPA:0091042) using the Human Protein Atlas via the gost function of the gprofiler2 R package (p-value = 0.004, ^58^). Interestingly, the cerebellum enriched transcripts showed allele specific changes amongst the *EXOSC3* p.G191 variants with Synaptotagmin-11 (SYT11) and UNC80 also showing gene dosage related changes (**Fig. 1C and Fig. S1E**). The RNA exosome complex has also been shown to degrade RNA transcripts with specific regulatory elements^46^. In these data, AU-rich elements in transcripts in DATs were significantly enriched for transcripts containing AU-rich elements (AREs) through RNA element enrichment analysis, suggesting a disrupted ability of the RNA exosome to regulate the decay of ARE-containing RNAs in G191D variants (**Fig. 1D**).

**Figure 1:**
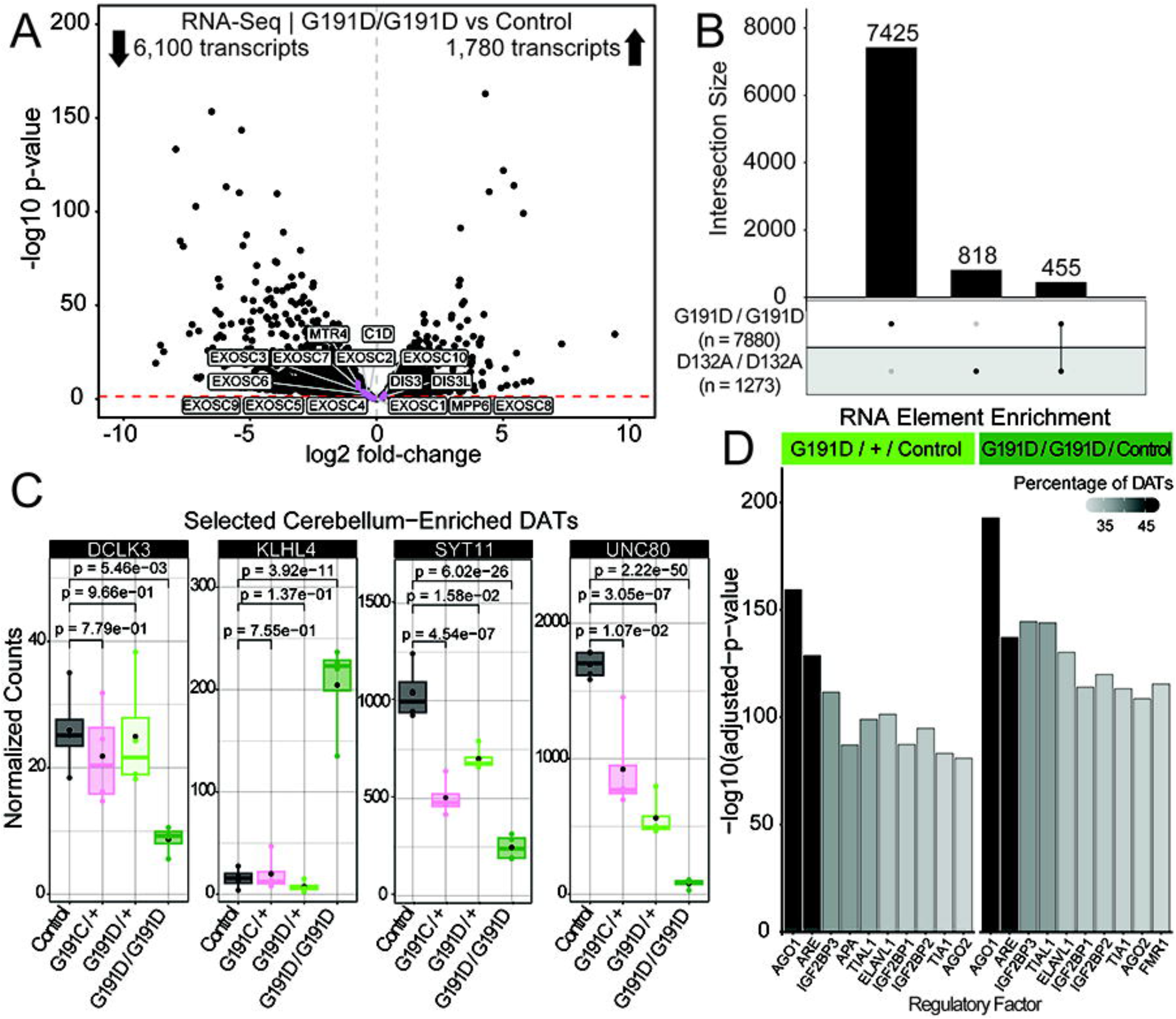
EXOSC3 G191D/G191D variant cells lines show extensive changes in transcript abundance relative to EXOSC3 D132A/D132A cell lines. **A.** Volcano plot of ultra-deep total RNA sequencing results from EXOSC3 G191D/G191D cells. Each dot represents a specific transcript across n=4 replicates. Transcripts for specific RNA exosome subunits are highlighted. The red dashed line indicates an adjusted p-value of less than or equal to 0.05. **B.** Upset plot illustrating the total number of differentially abundant transcripts (DATs) for each dataset and their overlap. The bars represent total number of DATs per category with their respective dataset presence indicated below by a black filled in circle. **C.** Normalized transcript expression graphs for a selection of cerebellum enriched DATs identified through pathway enrichment analysis. Each dot within the bar graph represents the value from a single biological replicate (n=4) with the bar representing the average abundance and the lines representing the standard deviation. **D.** RNA element enrichment analysis performed using AURA. Percentage of total DATs in each category is represented by the shading scale with the -log10 adjusted p-value for the analysis show on the y-axis.

Prior studies to characterize the role of EXOSC3 in transcriptome regulation have focused on the use of knockdown systems such as siRNA. Since PCH1B variants have been described as having loss of function in various model systems, it could be expected that the overlap between DATs in siRNA knockdown of EXOSC3 and in EXOSC3 PCH1B-related variants would be substantial. To determine if this is the case, we obtained the EXOSC3 (also known as RRP40) siRNA knockdown studies PRJNA292827 and PRJNA328219 from GEO and performed differential RNA-Seq analysis as described for our current studies to perform comparative analysis^59,60^. Following these analyses, we separated RNAs into biotype categories according to their ENSEMBL classification and plotted the DATs for each EXOSC3 G191 variant (**Fig. 2A**) and EXOSC3 siRNA knockdown dataset (**Fig. S2A**). Strikingly, EXOSC3 G191D/G191D cells displayed a median reduction in log2-transformed transcript fold change across all RNA classes when compared to the EXOSC3 G191 heterozygous cell lines (**Fig. 2A**). Conversely, the siRNA knockdown datasets reveal that siRNA knockdown of EXOSC3 leads to increased abundance of miRNAs, snoRNAs, lncRNAs, and snRNAs that was not observed in any of the EXOSC3 G191 variants. These data suggest that siRNA knockdown of EXOSC3 is not an accurate representation of the changes that occur in PCH1B variant expressing cells. Since these siRNA exposures are transient in nature, it is also possible that the G191D stable cell lines represent changes associated with system reequilibration associated with the genetic change in EXOSC3, as may occur in PCH1B variant carriers. Since prior work has shown that promoter associated transcripts (PROMPTs) significantly increase with EXOSC3 knockdown, we also performed genome wide averaging analysis for the level of PROMPTs in our EXOSC3 G191 cell lines relative to paired controls (**Fig. S2B**). This analysis revealed that PROMPTs are also not significantly increased in EXOSC3 G191 cell lines and may in fact decrease on average relative to control cell levels.

**Figure 2:**
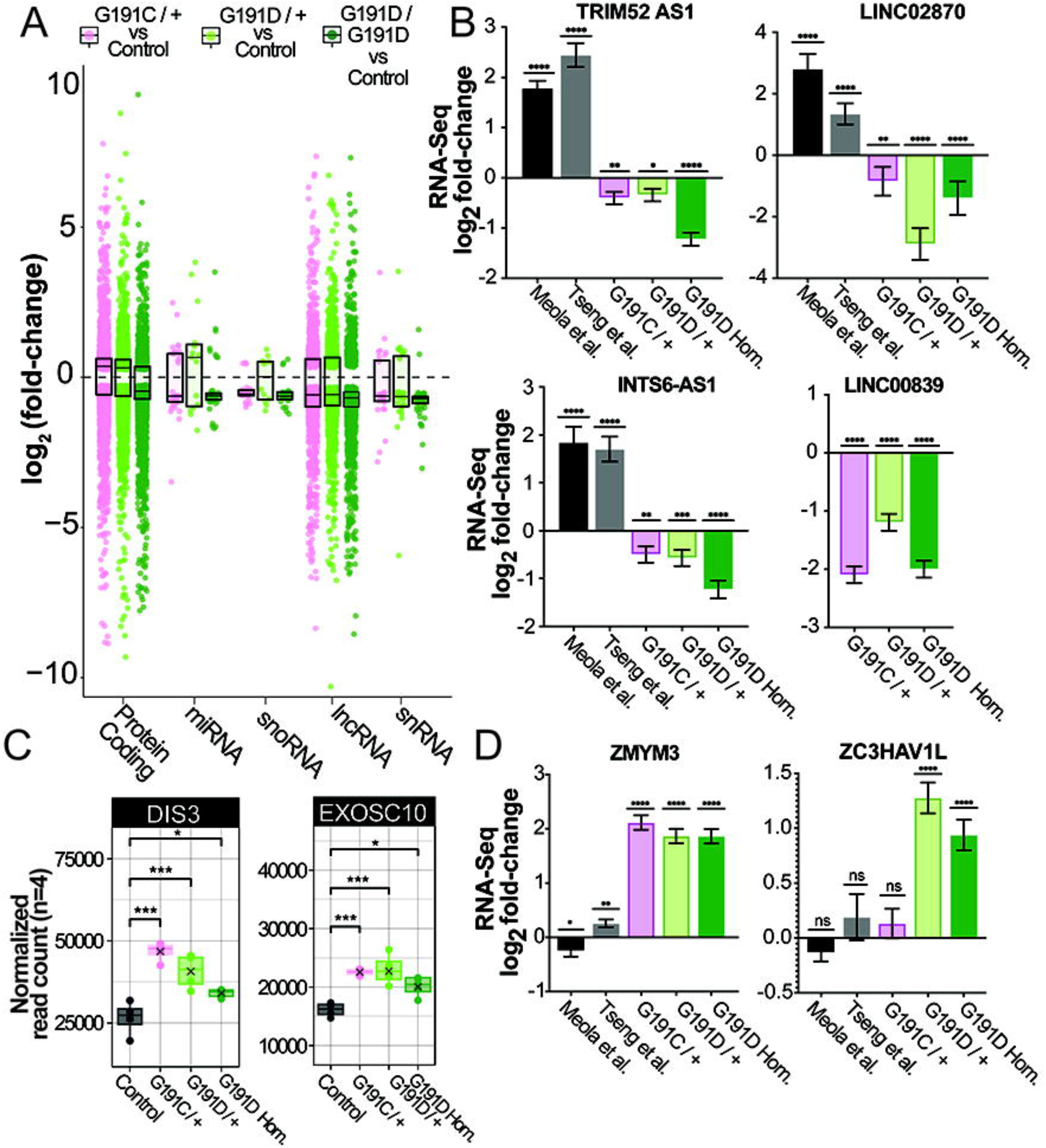
EXOSC3 G191 variant transcriptomes show distinct differences from siRNA EXOSC3 knockdown datasets. **A.** Transcript biotype analysis with the average DATs fold-change for each biotype shown as dots colored by EXOSC3 G191 variant genotype as indicated in the legend above the graph. Box plots show the median as a central line and the box shows the boundaries for the upper 75%tile or lower 25%tile of the datasets. **B.** Highlighted noncoding DATs from EXOSC G191 variant datasets or Meola, et al. / Tseng, et. al EXOSC3 siRNA knockdown datasets. Graphs show the average transcript abundance with the standard deviation (n<u>>3</u>) with transcript names indicated at the top of each graph. **C.** Normalized transcript expression graphs for 3’-5’ exonucleases DIS3 and EXOSC10. Each dot within the bar graph represents the value from a single biological replicate (n=4) with the bar representing the average and lines representing standard deviation. **D.** Highlighted DATs for zinc-finger proteins from datasets as indicated. Graphs show the average transcript abundance with the standard deviation (n<u>>3</u>) with transcript names indicated at the top of each graph. Significance values throughout the figure are *=p-valu**e** ≤ 0.05, **=p-value ≤ 0.01, ***=p-value ≤ 0.005, ****=p-value ≤ 0.001)

To further characterize these unexpected results, we performed overlap analysis of the significant DATs between each dataset (**Fig. S2C**). The EXOSC3 G191 dataset displayed lower total numbers of DATs when compared to the siRNA datasets with a high degree of DAT overlap identified between all three EXOSC3 G191 cell lines (**Fig. S2C**). A total of 363 DATs were identified across all five datasets and a selection of the expression profiles for these transcripts is shown in **Fig. 2B&D**. Many of the transcripts that were differentially expressed in all five datasets were noncoding RNAs, which are well established to be targets for the RNA exosome and can include PROMPTs, antisense (AS) transcripts, and long noncoding RNAs (lncRNAs) as shown in **Fig. 2B**^61–67^. A potential explanation for these findings was highlighted in **Fig. 1**, showing that some RNA exosome subunits are increased in transcript abundance in EXOSC3 G191 variant cell lines including the 3’-5’ exonucleases DIS3 and EXOSC10 and is shown for all replicates in **Fig. 2C**. Increased expression of the exonuclease subunits could compensate for reduced overall RNA exosome complex activity, especially in the case of DIS3 which has been shown to be functional independent of the fully assembled RNA exosome complex in vitro and recently in cells^68^. It has also been suggested that association of DIS3 and EXOSC10 with the RNA exosome decreases catalytic efficiency through allosteric effects^42,69^. The levels of the other core RNA exosome subunit transcripts did not significantly increase (**Fig. S3A**). Additional factors could also contribute to these effects either indirectly through regulation of transcript expression, splicing, and / or post-transcriptional RNA regulation. Of note for the EXOSC3 G191 datasets was the significant differential abundance of several zinc finger protein transcripts that were often not differentially expressed in the EXOSC3 siRNA datasets (**Fig. 2D**). In fact, pathway enrichment analysis of the differential transcripts in the EXOSC3 G191D/G191D vs. control dataset were strongly enriched for transcription regulators and metal ion binding proteins (**Fig. S4A**). In some cases, these nucleic acid regulatory proteins also showed an inverse response to EXOSC3 p.G191 variants than in the siRNA EXOSC3 knockdown studies (**Fig. S4B**).

### EXOSC3 G191 variants alter the balance of EXOSC3 isoforms through alternative splicing

Transcriptomic data also revealed thousands of splicing events unique to the G191 variants compared to control using *rMATs* software (**Fig. 3A**, ^70^). rMATs detects five specific splicing events: skipped Exon (SE) occurs when an exon is included in some transcripts but omitted in others; Alternative 5′ (A5SS) and 3′ (A3SS) Splice Sites involve different splice site selections, altering exon length; Mutually Exclusive Exons (MXE) ensure only one of two exons is included; Retained Intron (RI) occurs when an intron remains in the mature transcript, potentially affecting translation^70–72^. All findings from these alternative splicing analyses are summarized in **Fig. 3A** with quantitative data from *rMATs*. Among these splicing phenomenon, the *EXOSC3* transcript itself was significantly affected, with G191 variants promoting a switch to an EXOSC3 isoform with exon 3 being spliced out^73^. This change in EXOSC3 splicing was highly significant as only ∼50% of the transcripts exhibited a splice-in of Exon 3 in EXOSC3 G191D/G191D compared to control (**Fig. 3B**). Sashimi plots showing the average read counts for each genotype are shown at EXOSC3 which is transcribed from the bottom strand from right to left (**Fig. 3C**). These data show decreases in exon 3-related junction usage and a corresponding increase in exon 2 to exon 4 junction usage (**Fig. 3C**). Corresponding with genotype dosage, EXOSC3 G191D/+ heterozygous variants showed ∼25% of the transcripts as losing Exon 3 (**Fig. 3A**). This isoform switch likely represents a critical alteration in the regulation of *EXOSC3* transcripts in G191 variants, further implicating the variant in disrupting normal RNA homeostasis.

**Figure 3:**
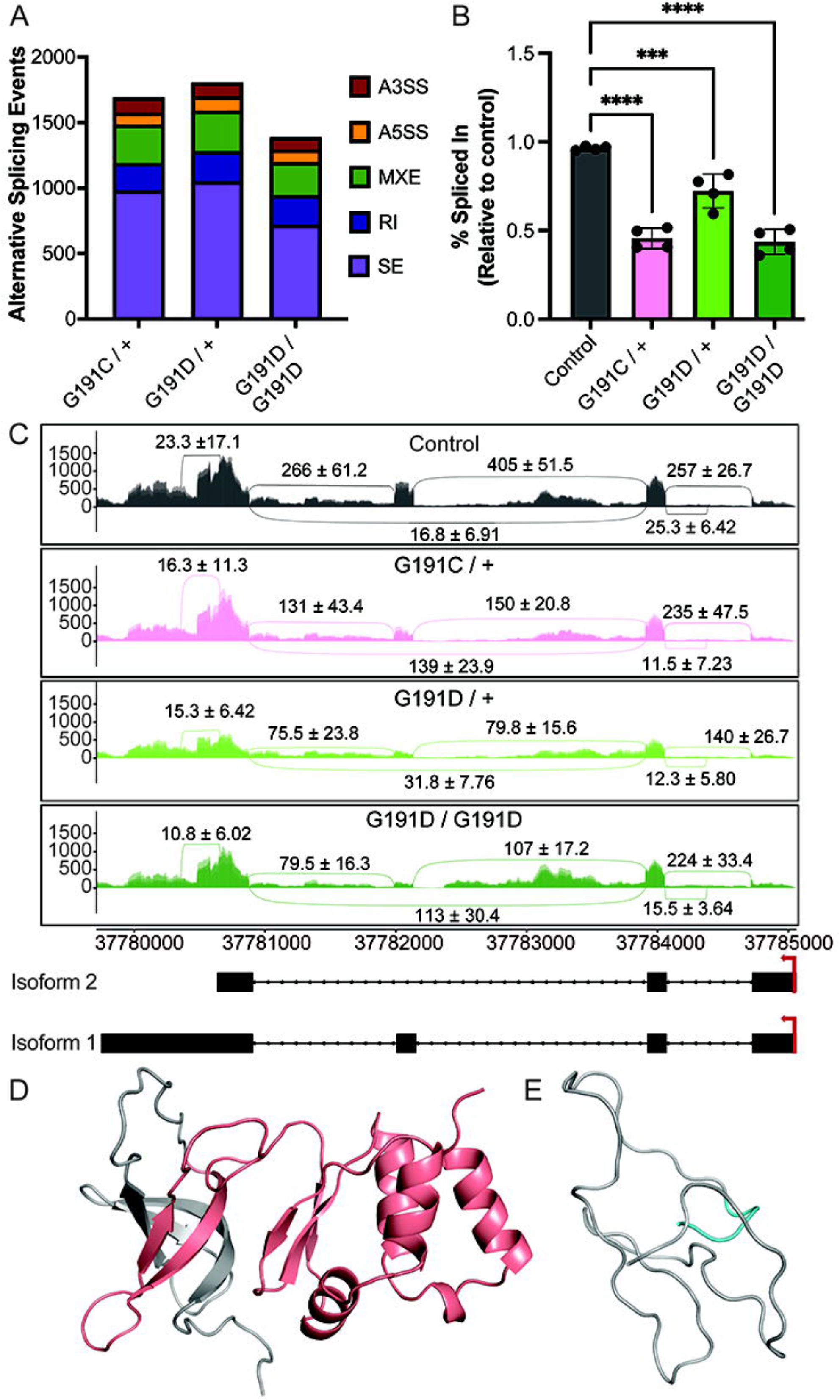
G191 variants cause alternative splicing of EXOSC3 transcripts. **A.** Summary graph of the significant alternative splicing events identified in each of the EXOSC G191 variant cell lines relative to controls. Summarized data includes alternative 5′- (A5SS) or alternative 3′- (A3SS) splice sites, mutually exclusive exons (MXE), retained introns (RI), and skipped exons (SE) and are color coded based on the legend to the right. **B.** Analysis of the percentage of exon 3 spliced-in for EXOSC3 in each genotype, as indicated at the bottom of the graph. Data are shown as average values plus or minus standard deviation across 4 replicates. Significance values are shown as ***=p-value ≤ 0.005 or ****=p-value ≤ 0.001. **C.** Sashimi plot and gene model annotation for EXOSC3 that contains two isoforms differing by inclusion / exclusion of exon 3 as shown in the annotations at the bottom of the figure. Genomic reads are converted into read densities per-base shown on the y-axis; junction reads are plotted as arcs, with the splice junction read counts indicated as averages with standard deviation (n=4) for each genotype as indicated. Snapshots from molecular dynamics simulations of the full length human EXOSC3 protein (**D**) or the shorter proteoform (**E**) that replaces the last 117 residues of EXOSC3 proteoform 1 with six different amino acid residues.

Considering that the alternative splicing event would generate a significant increase in the short transcript isoform for EXOSC3, we performed molecular dynamics simulations for each isoform to model how they might affect EXOSC3 protein structure and function. For these studies, the structure of the EXOSC3 subunit was obtained from a previously published RNA exosome structural model (PDB: 6D6Q)^52^. We first explored the alternative splicing event that replaces the last 117 residues of EXOSC3 with six different residues, leading to the loss of the entire C-terminal KH domain as well as the β4 and β5 strands of the S1 β-barrel. The effects of G191C and G191D-induced alternative splicing on EXOSC3 were examined by removing all the residues after 164 and using PyMOL’s mutagenesis tool to mutate residues 159-164 from VGDLIY to AISSRL^74^. In the absence of these structural elements, the S1 domain loses its integrity and, in our model, undergoes complete unfolding / destabilization, as observed in representative MD simulations (compare **Figs. 3D & E & S5 A & B**).These findings were supported by a lack of detection of the shorter isoform of EXOSC3 in any of our quantitative proteomics studies (discussed below) and an inability to detect the shorter isoform of EXOSC3 by western blotting using an N-terminal specific EXOSC3 antibody with beta-actin as a loading control (**Fig. S5C**).

Our data indicates that only the full length proteoforms of EXOSC3 containing G191 missense variants are stably expressed in G191D/G191D cells. Therefore, molecular dynamics (MD) simulations were performed for full length EXOSC3 G191, G191D, and G191C containing missense variants as either unbound EXOSC3 or RNA exosome-bound states. Upon analysis of the MD trajectories, the S1 domain’s native beta-motif-loop structure (β_5L_-L_5_-β_L5_) was observed to be disrupted by the two G191 variants. In native EXOSC3, the β_L5_ strand forms three hydrogen bonds with the adjacent β_4_ strand: G191 N – Y164 O, G191 O – Y164 N, and I193 N – L162 O, referred to as d_1_, d_2_, and d_3_, respectively (**Fig. 4A-C**). These hydrogen bonds are consistently intact in both complex and unbound simulations of the native EXOSC3 protein (**Fig. 4A**). However, G191D and G191C variants disrupt these three hydrogen bonds to varying degrees, with G191D disrupting them to a greater extent than G191C (compare **Fig. 4B to 4C**). Disruptions result from steric and electrostatic clashes introduced by the variant sidechains, as well as unfavorable dihedral angles sampled by non-Gly residues. For example, in G191C, d_1_ breaks due to its improper orientation toward β_4_. In the RNA exosome complex, d_1_, d_2_, and d_3_ interactions persist due to structural constraints of the assembled complex, but are weaker with elongated distances (∼3-4 Å). Collectively, the MD simulations show that neither G191C nor G191D adopt a single fixed conformation. Instead, G191C alternates between states where only d_1_ or all three hydrogen bonds are broken, while G191D exhibits more frequent and pronounced destabilization of all three β_L5_-β_4_ interactions. The introduction of non-Gly side chains also leads to hydrophobic packing disruptions around the 191 residue (compare **Fig. 4E&F to 4D**). In native G191 EXOSC3, V192 packs against hydrophobic residues within the interior of the S1 β-barrel (orange surface), M190 packs into the hydrophobic surface between the S1 β-barrel and the Kh domain (blue surface), and G191 associates tightly with β_4_ (**Fig. 4D**). In G191D, the hydrophobic contacts by V192 and G191D break (**Fig. 4E**) while in G191C, the G191C/β_4_ association is partially disrupted and M190 hydrophobic packing is sometimes lost (**Fig. 4E**). For both variants, these structural changes are observed to alter the behavior of the adjacent flexible L_5_ loop (compare **Fig. 4E&F to 4D**). To quantify β_L5_ conformational changes, we computed per-residue average backbone root mean square deviations (Δ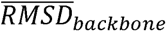) relative to the native G191 for both the bound (**Fig. 4G**) and unbound EXOSC3 (**Fig. 4H**). Both G191D and G191C exhibited increased Δ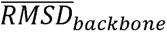i in β_L5_ and the L_5_ C-terminal region (residues 187-190), with more pronounced effects in unbound EXOSC3. Due to its greater frequency of detachment, G191D displayed higher Δ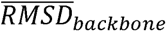 than G191C, reinforcing the stronger impact of its structural destabilization. λ-dynamics (λD) simulations were next performed to investigate 191 variant effects on EXOSC3’s folding stability (ΔΔ*G*_fold_) and binding affinity to the full RNA exosome complex (ΔΔ*G*_bind_, **Fig. 4I**)^75–81^. This investigation revealed that while G191D alone altered EXOSC3’s ΔΔ*G*_bind_ by slightly less than 1 kcal/mol, G191C displayed minimal impact on binding. However, both EXOSC3 variants, plus an extra G191A variant, induced large penalties to EXOSC3 folding with computed ΔΔ*G*_fold_ of ∼4-6 kcal/mol. Taken together, these results suggest that destabilization of EXOSC3 folding may have a larger contribution than EXOSC3’s RNA exosome binding affinity to G191 variant-induced PCH1b disease phenotypes (**Fig. 4I**). This analysis excludes possible reorganizational free energy penalties that may be needed to fold variant EXOSC3s into a suitable binding conformation, which would likely impact the true ΔΔ*G*_bind_. Notably, the lesser thermodynamic severity of the G191C variant compared to G191D correlates well with less severe clinical outcomes observed in G191C homozygous patients (previously classified with complicated hereditary spastic paraplegia) compared to G191D homozygous patients (classified with moderate PCH1b) ^23,53,54^.

**Figure 4:**
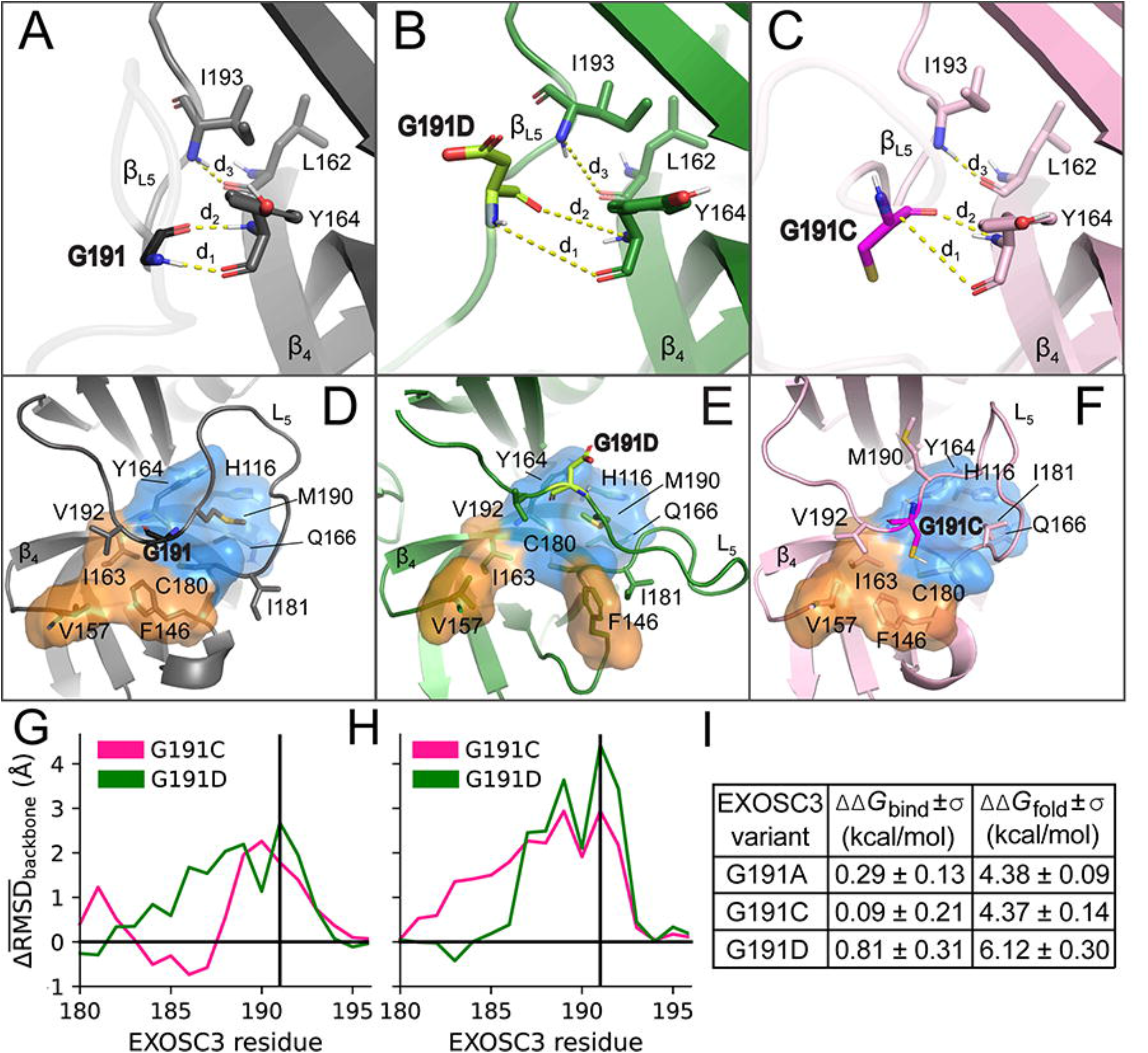
Molecular modeling of EXOSC3 G191C and G191D variant proteoforms. **A.** Molecular dynamics structures reveal key hydrogen bonds linking β_L5_ and β_4_ strands (d_1_: 191 N-Y164 O, d_2_: 191 O-Y164 N, d_3_: I193 N-L162 O) in G191 (β_L5_ tethered). **B.** same regions in EXOSC3 G191D. C. Same regions in EXOSC3 G191C. (**D-F**) G191C and G191D variant disrupt hydrophobic interactions near M190 and V192, leading to β_L5_ destabilization. In the EXOSC3 reference state (**D**), M190 binds a hydrophobic pocket (H116, Y164, Q165, C180), G191 interacts with C180, I163, and F146, and V192 contacts I163, F146, and V157. These interactions weaken in G191D/C, causing V192 to lose contact and M190 to flip out, with I181 filling the space (**E-F**). (**G-H**) Δ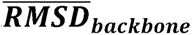 of L_5_, β_L5_, and S1/KH linker shows structural deviations, with residue 191 marked for (G) EXOSC3 is bound to the RNA exosome and (H) EXOSC3 in isolation. (I) ΔΔ*G*_bind_ and ΔΔ*G_f_*_old_ for EXOSC3 G191 variants G191A, G191C, and G191D computed via λ-dynamics with analytic correction. Data shown are average ΔΔ*G*_bind_ and error (± σ) values.

### EXOSC3 G191 variants alter system-wide RNA processing enzyme proteostasis

Given the effects of G191 variants on the RNA exosome complex, as measured by molecular dynamics simulations, and their impact on transcriptome abundance levels, we hypothesized that these single amino acid changes in EXOSC3 p.G191 would also lead to proteome-wide perturbations. To test this, quantitative mass spectrometry-based proteomics was performed to measure global protein abundance on an Orbitrap Eclipse (**Fig. 5A**). Of the 7,784 proteins detected, we quantified 6,856 including all RNA exosome subunits and a number of relevant cofactors. Pearson correlation coefficients between the biological replicate measurements for the proteomics datasets ranged from 0.980 - 0.991 for EXOSC3 G191D/G191D homozygous cell lines and from 0.965 - 0.993 for the isogenic controls (**Fig. S6**). Correlation between the mRNAs for the paired RNA-Seq relative to the global proteomics datasets showed lower positive Pearson correlation values ranging from 0.333 – 0.378 for all genotypes tested which is consistent with prior studies on transcriptome / proteome correlation ^82–84^.

**Figure 5:**
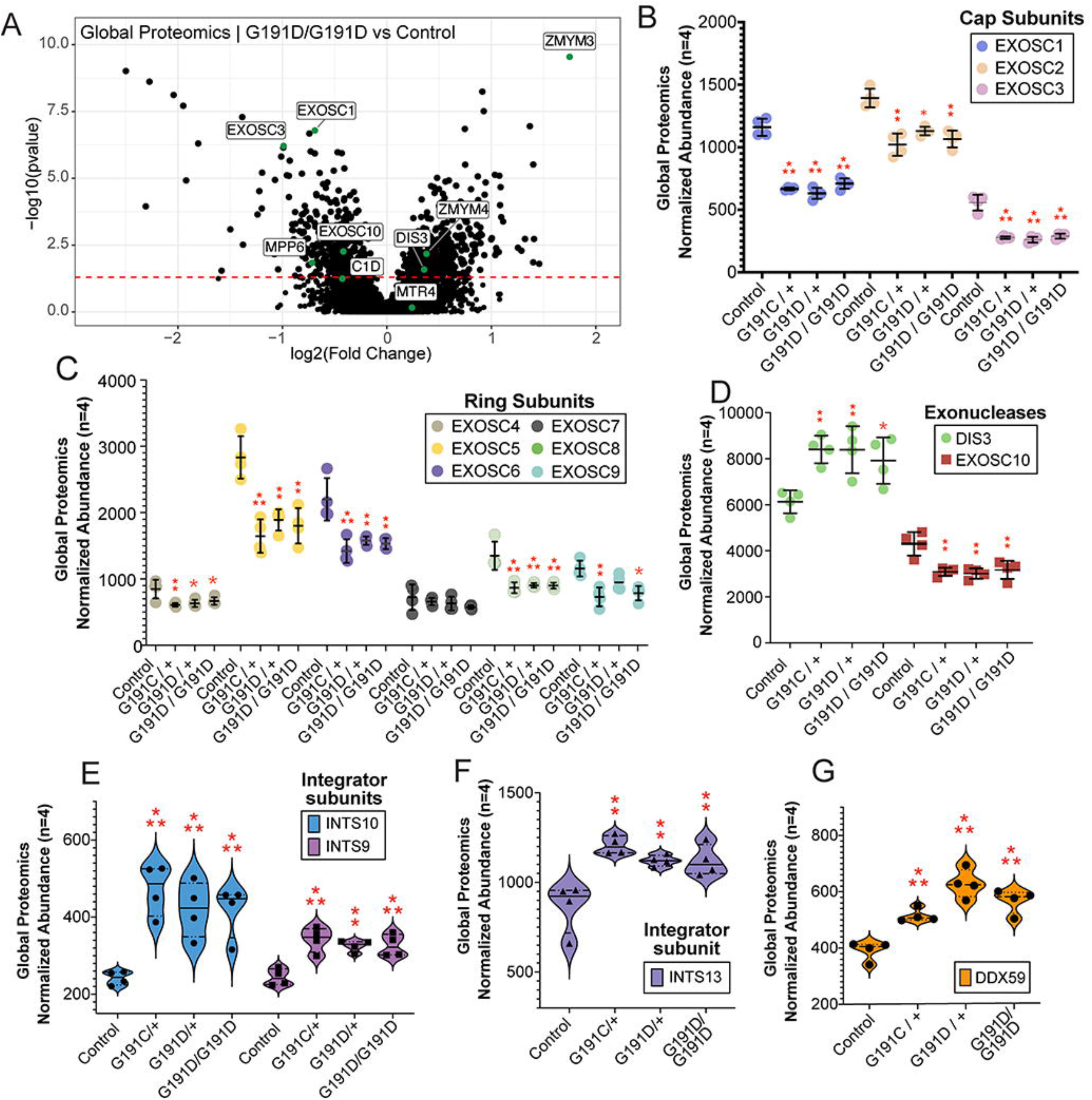
Expression of EXOSC G191 variants alters key RNA processing protein abundance. **A.** Volcano plot of global proteomics studies in EXOSC3 G191D/G191D homozygous variant cell lines versus controls (n=4). Protein abundance was quantified using total TMT reporter ion intensity and analyzed with Proteome Discoverer 2.5. Differentially abundant proteins (DAPs) were identified using p-value < 0.05. The volcano plot displays log2 (fold change) versus -log10(p-value), highlighting proteins of interest in the RNA exosome complex and the zinc finger proteins ZMYM3 & 4 with green dots. (**B-D**) Normalized abundance graphs for subunits of the RNA exosome for the cap subunits (**B**), ring subunits (**C**), and exonuclease subunits (**D**). Data is presented with a dot for each biological replicate sample (n=4) with the average represented by a central line plus or minus the standard deviation. (**E-F**) Violin plots illustrating the abundance values for RNA processing proteins including various Integrator subunits (**E & F**) and DEAD-box helicase DDX59 (G). Significance values throughout the figure are *=p-value ≤ 0.05, **=p-value ≤ 0.01, ***=p-value ≤ 0.005.

Global proteomic analysis revealed significant alterations in the abundances of several RNA exosome subunits, including EXOSC1, EXOSC2, EXOSC3, EXOSC4, EXOSC5, EXOSC6, EXOSC8, EXOSC9, EXOSC10, and DIS3 (**Fig. 5A, Fig. S7A & B**). Across the four biological replicates, EXOSC3 showed a 50% reduction in protein abundance across all three genotypes, with statistically significant fold changes of 0.475 (G191C/+), 0.45 (G191D/+), and 0.504 (G191D/G191D) relative to control (**Fig. 5B**). When comparing the protein level and RNA level of the RNA exosome cap subunits, we saw RNA levels of RNA exosome subunits either decreasing or not changing (**Fig. S3**). *EXOSC3* transcript levels have a fold decrease of 0.765 in G191D/G191D homozygous cells, suggesting that the EXOSC3 protein levels are relatively lower than its decrease in transcript levels, possibly as a consequence of protein destabilization and related turnover (described in **Fig. 4**). In addition, we observed reduced protein level abundance of the cap subunits EXOSC1 and EXOSC2 in cells carrying at least one variant allele (**Fig. 5B**). At the RNA level, EXOSC1 levels did not significantly change in any of the genotypes relative to controls. Reductions in RNA exosome ring subunit protein levels were also observed across all EXOSC3 G191 variant samples (**Fig. 5C**). Interestingly, despite the general decrease in most RNA exosome subunits in protein abundance, the 3’-5’ exonuclease subunit DIS3 was upregulated by approximately 1.25-fold relative to the control cell line (**Fig. 5D**). Both 3’-5’ exonucleases in the complex (DIS3 & EXOSC10) were upregulated at the mRNA level (**Fig. 2C**), however EXOSC10 protein levels were significantly decreased across all G191 variant expressing cell lines. Protein levels of some other RNA processing factors implicated in RNA processing and RNA degradation were also upregulated at the protein level including the Integrator subunits INTS9, INTS10, and INTS13 in all EXOSC3 G191 cell lines relative to isogenic controls (**Fig. 5E & F**). The Integrator complex is known to play a role in premature RNA Polymerase II transcription termination of mRNAs in addition to processing of other small RNAs through mechanisms that require the RNA exosome^85–87^. The DEAD-box helicase DDX59 is also significantly increased in EXOSC3 G191 cell lines (**Fig. 5G**). DDX59 has recently been implicated in the degradation of the nuclear noncoding RNA MALAT1 whose levels are also regulated by the RNA exosome^88^.

### G191 variants affect RNA exosome cap subunit and MPP6 stability and interactions

Our analysis revealed a widespread decrease in the abundance of most RNA exosome subunits in cells expressing G191 variants. To experimentally test whether these protein-level changes were associated with alterations in protein complex integrity and/or protein stability changes, we assessed proteome-wide thermal stability using the Protein Integral Solubility Assay (PISA) in all EXOSC3 p.G191 and control lines (**Fig. 6A**, **Fig. S8A & B**). By PISA, significant destabilization of EXOSC1, EXOSC3, and MPP6 was observed in G191D/G191D homozygous cells (**Fig. 6A**). These findings are consistent with our MD and λD predictions for the EXOSC3 G191 missense variants as shown in **Fig. 4**. In the G191D/+ and G191C/+ heterozygous cell lines, significant protein destabilization was also observed for EXOSC1 and MPP6 (**Fig. S8A & B**). For EXOSC3, thermal stability was significantly reduced in G191D/G191D homozygous cells, whereas G191C/+ and G191D/+ heterozygous variants exhibited more subtle changes suggesting that the presence of reference EXOSC3 in G191 variant containing heterozygous cells can partially mitigate the average change in EXOSC3 thermal destabilization, as would be expected. These findings suggest that G191 variants primarily impact the thermal stability of EXOSC3 and its direct binding partners, potentially disrupting interactions within the RNA exosome complex (**Fig. 6A**). In contrast, while other subunits of the RNA exosome exhibited reduced abundance by global proteomics, their thermal stability remained largely unaffected, indicating that their depletion may occur through mechanisms independent of protein destabilization, such as altered synthesis or degradation pathways that could include orphan protein decay^89^.

**Figure 6:**
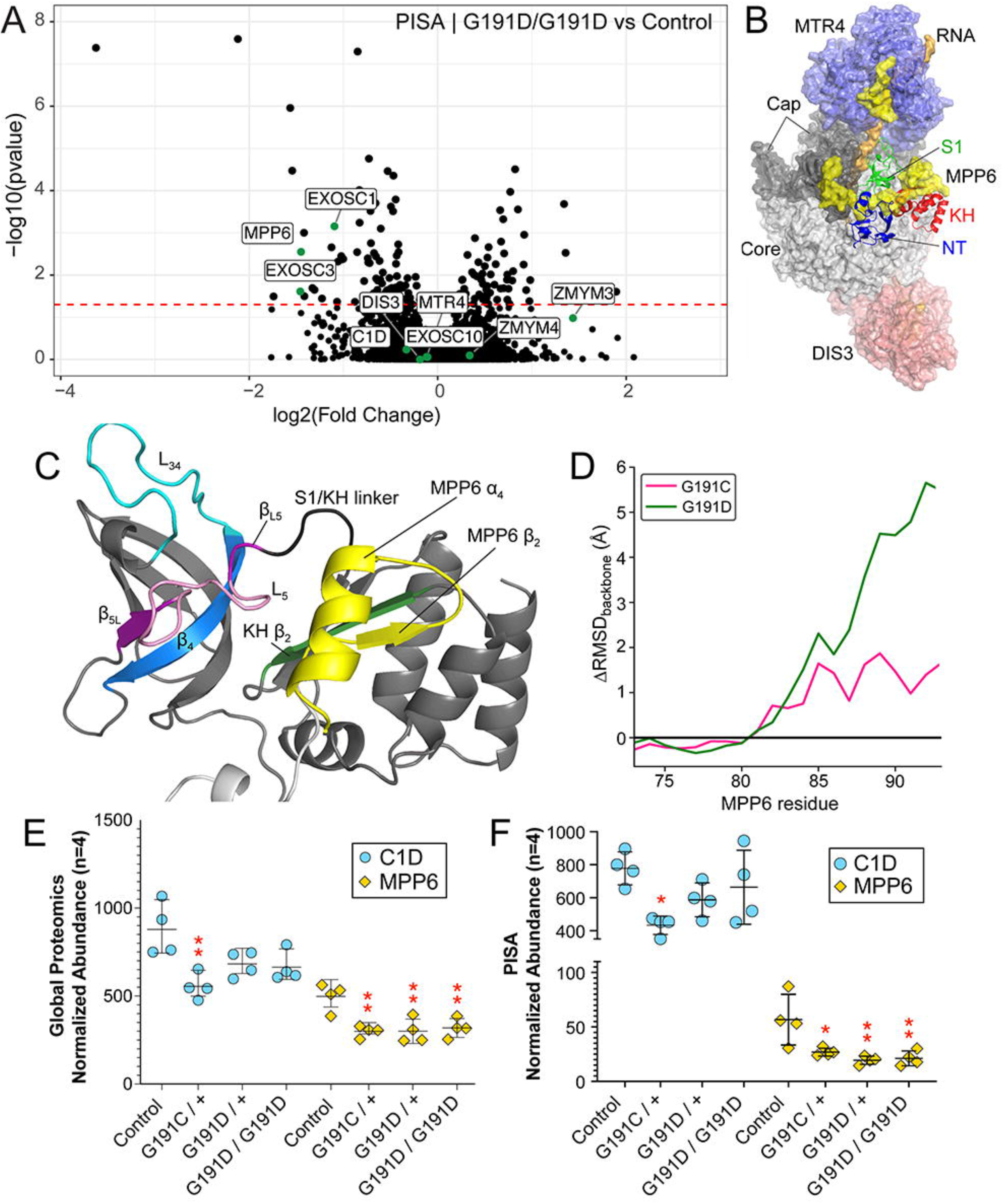
Protein thermal stability analysis reveals destabilization of some RNA exosome cap subunits and the RNA binding protein MPP6. **A.** Volcano plot of PISA proteomics studies in EXOSC3 G191D/G191D homozygous variant cell lines versus controls (n=4). Protein abundance was quantified using total TMT reporter ion intensity and analyzed with Proteome Discoverer 2.5. Differentially abundant proteins (DAPs) were identified using p-value < 0.05. The volcano plot displays log2 (fold change) versus -log10(p-value), highlighting proteins of interest in the RNA exosome complex and the zinc finger proteins ZMYM3 & 4 with green dots. B. Surface representation of the RNA exosome structure PDB: 6D6Q from Weick et al., 2018 with MTR4 shown in blue, DIS3 shown in red, RNA exosome cap subunits shown as dark gray, the core subunits shown in light gray, MPP6 highlighted in yellow, and EXOSC3 shown as a ribbon diagram to show EXOSC3 proximity to MPP6. C. Ribbon diagram of the EXOSC3 and MPP6 interface from PDB: 6D6Q. D. Δ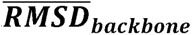 of MPP6 residues 73-92. (E-F) Normalized abundance graphs for MPP6 and C1D from either global proteomics (E) or PISA (F) for all genotypes as described under the figure. Data is presented with a dot for each biological replicate sample (n=4) with the average represented by a central line plus or minus the standard deviation. Significance values are *=p-value ≤ 0.05, **=p-value ≤ 0.01.

Previously reported structural analysis of the RNA exosome containing both the RNA helicase MTR4 and the RNA binding protein MPP6 revealed the close association of MPP6 with EXOSC3 in the cryo-EM human complex (**Fig. 6B&C,** ^52^). MD analysis suggested that, as a result of EXOSC3 G191 missense variants, the L_5_, β_L5_, and the S1/KH linker structures could form into a single unstructured loop (**Fig. 4D-F**), disrupting normal interactions with MPP6 in the RNA exosome complex and the KH domain in EXOSC3. However, structural constraints within the RNA exosome complex could partially mitigate these effects. To investigate how these G191 variant conformational changes affect MPP6 structure and binding, we calculated a per-residue 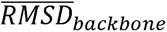 for MPP6 residues 74-93 (**Fig. 6D**). Residue 93 is the last MPP6 residue included in the cryo-EM RNA exosome model. However, human MPP6 contains more than 60 additional C-terminal residues that are missing from all current cryo-EM structures. When calculating the per-residue 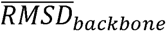 between the G191 variants, the G191C-bound MPP6 α_2_ displayed a per-residue 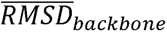 of 1-2 Å, and the G191D produced a much larger effect of 2-6 Å (**Fig. 6D**). While a more complete structure of MPP6 could attenuate the changes seen in MPP6 α_2_, the MPP6 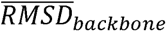 results suggest G191 missense variants increase the 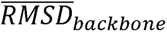 of α_2_ within the complete MPP6, with G191D producing a larger effect than G191C, likely impacting the stability of MPP6 – EXOSC3 interactions. MPP6 has been linked to activating and recruiting the MTR4 helicase, which could be negatively impacted by the disruption of MPP6’s α_2_ structure.

Prior work has shown that MPP6 associates with the nucleic acid-binding protein C1D and MTR4 in protein-protein interaction studies^90^. Since our MD analysis suggests that MPP6 binding interactions could be altered by the EXOSC3 G191 variants, we investigated specific PISA changes for these three RNA exosome regulatory factors. We did not observe changes in either MTR4 protein abundance or MTR4 protein stability, suggesting that MTR4 structure is not significantly affected (**Fig. S8C & D**). However, we did observe significant changes in protein stability for both MPP6 and C1D. MPP6 significantly decreased in both global abundance and PISA-measured thermal stability in all EXOSC3 G191 datasets (**Fig. 6A, E & F**). Conversely, significant decreases in abundance and thermal stability of C1D only occurred in the EXOSC3 G191C/+ datasets suggesting that the G191C variant may cause allele-specific changes in C1D association and/or recruitment to the RNA exosome (**Fig. 6E&F, Figs. S7A & S8A**).

### Recalibration of specific RNA processing factors occurs with EXOSC3 overexpression

Our findings show that EXOSC3 protein abundance and thermal stability are significantly altered in EXOSC3 variants through a combination of changes. First, ultra-deep RNA Seq revealed that EXOSC3 G191 variant cells have increased relative production of the second EXOSC3 transcript isoform that lacks exon 3 leading to a destabilized version of the protein (**Fig. 3**). As a consequence, total EXOSC3 protein abundance is significantly decreased as is the abundance of eight of the nine core subunits of the RNA exosome (**Fig. 5**). Overall EXOSC3 levels were also significantly decreased in EXOSC G191 variant heterozygous cell lines (**Fig. S7A & B**). Furthermore, PISA thermal stability analysis revealed that the full length EXOSC3 in G191D/G191D homozygous cells was significantly thermally destabilized relative to reference controls **(Fig. 6)**. EXOSC3 G191 variant heterozygous cell lines, however, did not show significant thermal destabilization of EXOSC3 (**Fig. S8A & B**). Considering that carriers of single copy EXOSC3 variants do not present with PCH1B, these findings suggest that either the overall abundance of EXOSC3 and/or the protein instability are the likely cause of human neurodegenerative disease. Using our isogenic cell models, we can address these possibilities by testing for genetic rescue through overexpression of either reference EXOSC3 or G191D EXOSC3 that we have shown has significant protein destabilization. For these experiments, we have produced EXOSC3-HaloTag overexpression plasmids from cDNA so alternative splicing of the rescue transcripts will not be a consideration. In brief, control or EXOSC3 G191D/G191D cells were transiently transfected with no DNA (mock transfection), WT EXOSC3, or G191D EXOSC3 and then the plasmid exposed cells were FACS sorted to isolate positive transfectants. Three biological replicates per condition were used for global proteomics analysis to assess the system-wide changes that are EXOSC3 dependent.

As shown in **Figure 7A**, both EXOSC3 WT and G191D expressing plasmids led to significant overexpression of EXOSC3 compared to mock transfected cells, although the expression level of EXOSC3 G191D was significantly lower in control cells than WT (p-value=0.05). Since both EXOSC1 and MPP6 showed significant reductions in both protein abundance and thermal stability in our EXOSC3 G191D/G191D variant lines relative to control (Figs. 5A & 6A), we assessed if overexpression of WT or G191D EXOSC3 could rescue their protein levels. Although neither EXOSC1 or MPP6 levels were significantly altered with EXOSC3 overexpression in the control cell line, both proteins showed significant increases in EXOSC3 G191D/G191D cells with overexpression of either WT or G191D EXOSC3 (**Fig. 7B & C**). Significant increases in core RNA exosome subunit EXOSC5 protein levels were also observed with WT EXOSC3 overexpression (**Fig. 7D**). Since we performed global proteomics analysis of this study, we were also able to identify other significantly different proteins that responded to EXOSC3 overexpression such as SKI2 and SKI3, subunits of the SKI complex that works with the RNA exosome in the cytoplasm to carry out mRNA decay and mRNA quality-control regulation^91,92^. SKI2, SKI3, and SKI8 were significantly increased in abundance in EXOSC3 G191D/G191D cell lines relative to controls in the mock transfected samples (**Fig. 7E & F**, p<0.001, p<0.001, and p<0.005 respectively). Upon EXOSC3 G191D overexpression, levels of SKI2 and SKI3 significantly decreased only in EXOSC3 G191D/G191D cell lines (**Fig. 7E & F**). The DEAD-box helicase DDX59 was significantly upregulated in EXOSC3 G191D/G191D cell lines relative to controls with mock transfection in this dataset in addition to the full EXOSC3 G191 variant analysis (p<0.001, Fig. 5G). However, DDX59 protein abundance levels did not significantly change with EXOSC3 WT or G191D overexpression.

**Figure 7:**
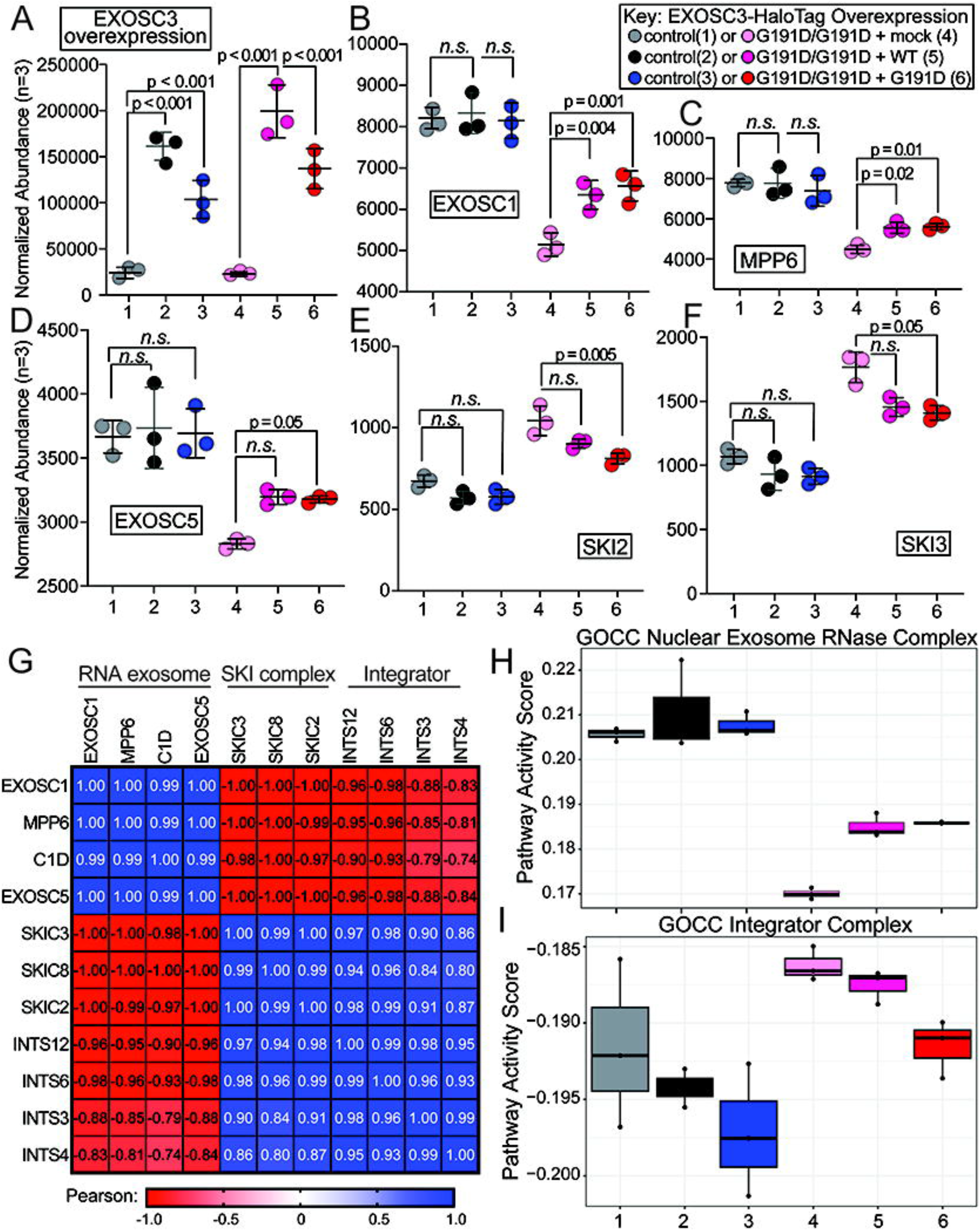
EXOSC3 protein levels are limiting for RNA exosome complex assembly in EXOSC3 G191D/G191D cell lines. **A.** Normalized abundance values for EXOSC3 in control or transient overexpression experiments color coded as shown in the key on the upper left. Data for this and subsequent graphs is presented with a dot for each biological replicate sample (n=4) with the average represented by a central line plus or minus the standard deviation. Significance values are given above the brackets showing the specific comparisons that were made. (B-F) Normalized abundance values for EXOSC1 (**B**), MPP6 (**C**), EXOSC5 (**D**), SKI2 (**E**), SKI3 (**F**) in the overexpression experiments. G. Pearson correlation matrix comparing the normalized abundance values for the proteins indicated at the top of the figure to the normalized abundance values for each protein indicated to the left of the figure. The Pearson correlation color scale is shown under the figure. (**H-I**) Pathway Activity Score analysis shown for the Nuclear RNA exosome (H) or the Integrator Complex (**I**).

Our global proteomics analysis in EXOSC3 G191 variant cells lines revealed that several RNA Processing enzymes including subunits of the Integrator complex and DDX59 were significantly increased in EXOSC3 variant cell lines compared to controls (**Fig. 5**). To determine if these proteome changes were responsive to EXOSC3 overexpression, we performed correlation analysis of the normalized protein abundance values for each factor of interest across all 18 samples in the overexpression dataset with a focus on the RNA exosome, the SKI complex, and the Integrator complex (**Fig. 7G**). RNA exosome related proteins EXOSC1, MPP6, C1D, and EXOSC5 showed high positive correlation (Pearson r = 0.99-1.00) with each other, suggesting that they all had similar abundance changes in response to EXOSC3 G191 status and overexpression changes (**Fig. 7G, upper left**). The protein abundance levels for subunits of the SKI complex (SKI2, SKI3, and SKI8) and the Integrator complex (INTS3, INTS4, INTS6, and INTS12) also showed high levels of correlation (Pearson r = 0.80-0.99) with each other across the EXOSC3 overexpression dataset (**Fig. 7G, lower right**). Strikingly, the protein abundance levels of the subunits of the RNA exosome complex showed a strong anti-correlation (Pearson r = -1.00 – -0.74) to the protein levels of the SKI and Integrator complexes (**Fig. 7G, lower left and upper right**). These data suggest that the levels of the SKI and Integrator complex subunits are highly responsive to EXOSC3 status across the overexpression dataset. Pathway activity score (PAS) analysis with a focus on Gene Ontology Cellular Components (GOCC) revealed similar results. Note that EXOSC3 protein abundances were removed for PAS so that exogenous EXOSC3 overexpression did not skew the enrichment findings. GOCC annotations for the Nuclear RNA exosome RNase Complex showed significant decreases in EXOSC3 G191D/G191D cell lines relative to controls with mock transfection (**Fig. 7H, compare sample 1 to 4**). The Nuclear RNA exosome PAS increased with overexpression of either WT or G191D EXOSC3 (**Fig. 7H, compare 4, 5, & 6**). Conversely, the GOCC for the Integrator complex increased when comparing EXOSC3 G191D/G191D cell lines relative to controls with mock transfection (**Fig. 7I, compare sample 1 to 4**). Integrator complex PAS decreased with overexpression of either WT or G191D EXOSC3 (**Fig. 7I, compare 4, 5, & 6**). Together, these findings show that EXOSC3 overexpression improves the levels of multiple subunits of the RNA exosome that are decreased in global abundance in EXOSC G191D/G191D cell lines. Additionally, RNA processing factors including subunits of the SKI and Integrator complexes are highly responsive to functional EXOSC3 levels as a possible compensation mechanism for reduced RNA exosome activity.

## Discussion

The RNA exosome is a central regulator of RNA homeostasis, contributing to RNA quality control, RNA processing, and RNA decay. Tissue specific diseases such as PCH1B are associated with cell or tissue-specific pathway disruption, suggesting that other human cells may be able to recalibrate or adjust to disease-associated variants. Our study investigates this through analysis of EXOSC3 variants that are associated with milder forms of PCH1B. The specific mechanisms that underlie allele-specific disease severity have not been investigated. Our findings identified several striking results, including that heterozygous EXOSC3 G191 variants appear to cause system disruption that is less severe than the homozygous EXOSC3 G191D/G191D cells but still extensive. Since heterozygous carriers of EXOSC3 G191 variants do not show symptoms of disease, this clearly suggests that cells are able to compensate for EXOSC3 functional disruption as we have observed in these studies. Our multiomics studies show that cellular compensation occurs at both the transcript and protein levels and involves adjustments within the RNA exosome complex itself and throughout a variety of other RNA processing and decay pathways.

Our analyses revealed significant and widespread changes in gene expression (DATs) and splicing profiles in EXOSC3 G191 variant cells. Interestingly, comparison to deep RNA sequencing studies performed in EXOSC3 D132A/D132A cells suggested that many of the changes are allele-specific with EXOSC3 G191D/G191D causing more extensive DATs. However, shared DATs included several RNAs that have known enrichment in the cerebellum as well as enrichment in transcripts with AU-rich elements, that could contribute to reduced neuronal health/survival in individuals with PCH1B in agreement with prior work^93,94^. Surprisingly, we also found that EXOSC3 G191D/G191D transcriptomes showed average decreases in the population of lncRNAs and other noncoding RNA species, in contrast to prior RNA-seq studies using transient siRNA-based knockdown of EXOSC3 where noncoding RNAs increase^59,60^. We propose that these changes are associated with increased production of the 3’-5’ exonuclease DIS3 in G191D/G191D EXOSC3 cells at both the transcript and protein levels. DIS3 is one of the catalytic subunits of the RNA exosome although it has also recently been shown to function independently of the RNA exosome complex in the cytoplasm^68^. These findings show that EXOSC3 siRNA knockdown does not accurately phenocopy transcriptome changes observed with EXOSC3 variants related to PCH1B. We also report the novel finding that EXOSC3 G191 transcripts show significant increases in exon 3 skipping, leading to production of a short transcript isoform whose translation would produce a highly unstable EXOSC3 proteoform. Through parallel studies of homozygous EXOSC3 D132A, G135R, and A139P sequence variants, we can report that increased exon 3 skipping is currently unique to EXOSC3 G191 variants. Together, these findings show dosage suppression of EXOSC3 G191 variant effects through DIS3 upregulation^95^. This is a unique finding since literature broadly has reported common co-regulation of genes in protein complexes at both transcription and translational levels^96–102^. These findings could potentially explain the observation of intrafamily variability of EXOSC3 G191 variant associated disease severity^54^.

In addition to upregulation of the 3’-5’ exonuclease DIS3 in G191D/G191D EXOSC3 cells, we also observe alterations in the balance of other transcription and RNA processing associated proteins. This included upregulation of a number of zinc-finger containing proteins such as ZMYM3, which was ∼4-fold induced at both the transcript and protein levels. ZMYM3 is a transcriptional regulator whose mutation is associated with neurodevelopmental delay^103^. ZMYM3 has been shown to interact with the RNA endonuclease RNAse H2, which resolves DNA-RNA hybrids^104^. Interestingly, RNAse H2 has been shown to act redundantly with the RNA-DNA helicase Senataxin, which recruits the RNA exosome to the genome through a direct interaction with EXOSC9^105,106^. Together, these findings suggest increased ZMYM3 levels could be associated with increased recruitment of RNAse H2 to the genome to potentially mitigate decreases in RNA exosome function. We also observed increased levels of several subunits of the RNA endonuclease complex Integrator at both transcript and proteins levels. The Integrator complex copurified with the RNA exosome in prior work showing the role of both Integrator and the RNA exosome in premature RNAPII transcription termination^87^. Upregulation of subunits of the Integrator complex could potentially increase the efficiency of RNA exosome recruitment when the complex is in limited abundance or possibly compensate for its decrease through Integrators endonuclease activity. The DEAD-box helicase DDX59, which has recently been implicated in noncoding RNA degradation along with multiple subunits of the RNA exosome, was also significantly upregulated at both transcript and protein levels^88^. The upregulation of several major cellular RNA degraders in EXOSC3 G191D/G191D cells could suggest a multi-target regulatory network for cellular compensation in response to decreased RNA exosome / degrader activity.

In this work, we have performed a multi-modal analysis of protein-level alterations caused by EXOSC3 G191 variants that included molecular dynamics, λ-dynamics, global proteomics abundance measurement, and thermal stability assessment of the proteome using PISA. This comprehensive analysis revealed that EXOSC3 has large predicted changes in protein folding ranging from ΔΔ*G*_fold_ of 4.37-6.12 kcal/mol, with the largest changes occurring due to the G191D sequence variant. These findings are supported by a significant decrease in EXOSC3 G191D/G191D protein abundance and thermal stability identified through global proteomics and PISA experiments. Although EXOSC3 protein level decreases were observed in the global proteomics experiments for the heterozygous G191 variants, thermal stability changes did not reach significance in the heterozygous cell lines, likely due to the presence of the reference EXOSC3 allele. Despite these observations for EXOSC3, significant thermal destabilization was observed in all genotypes for the RNA exosome cap subunit EXOSC1 and the RNA binding protein MPP6, both of which directly interact with EXOSC3. These findings are consistent with our molecular dynamics and λ-dynamics simulations showing that protein-protein interactions are weakened and/or disrupted locally in the RNA exosome complex because of the presence of the EXOSC3 G191 variant. Interestingly, all nine of the RNA exosome core components (collectively also known as Exo9) were significantly decreased in global abundance as was the 3’-5’ exonuclease EXOSC10 that associates with the cap subunits including EXOSC3. We conclude that the reduction of Exo9 and EXOSC10 protein abundance across all subunits is caused by orphan protein decay because of the increased production of the unstable, and likely unusable, short EXOSC3 proteoform^107–109^. Since DIS3 levels increase significantly in all EXOSC3 G191 variants with no concurrent change in DIS3 thermal stability, our findings also suggest that DIS3 does not require Exo9 to escape orphan protein decay, making it unique among all RNA exosome subunits.

Prior work has suggested that the cerebellum may have a high requirement for RNA exosome function such that loss of RNA exosome expression, function, and/or insufficient protein complex formation in patients with EXOSC3 pathogenic variants lead to pathology^110^. Together with our findings at the protein level, these data suggest that decreased levels of some subunits of the RNA exosome could be sufficient to cause PCH1B. Our overexpression experiments further showed that after 72 hours, expression of either reference or G191D EXOSC3 was sufficient to increase the abundance of other Exo9 subunits including EXOSC1 and EXOSC5 in addition to the regulatory protein MPP6. Surprisingly, we also found significant decreases in the abundance of SKI complex subunits because of EXOSC3 overexpression. Additional correlation studies showed that as RNA exosome subunit levels increase in EXOSC3 G191D/G191D cells with plasmid EXOSC3 overexpression, levels of the SKI complex and the Integrator complex decrease. These findings suggest that not only are other RNA processing factors being upregulated because of decreased RNA exosome function, but that those system wide changes can be reversed with RNA exosome rescue. All together, these finding suggest the use of an antisense oligonucleotide or small molecule could be a potential therapeutic intervention specifically for patients with *EXOSC3* p.G191 variants to reduce *EXOSC3* exon 3 skipping thereby increasing the total levels of EXOSC3 protein.

## Methods

### Cell Culturing & Cell Line Generation

The parental cell line used to create the variant cell lines is HEK293T purchased from American Type Culture Collection (ATCC). The HEK293T cell line were edited using homology direct repair. The cells were transfected with plasmids containing CRISPR/Cas9, gRNA that targeting EXOSC3-G191, and a DNA template that contained synonymous variant for AscI restriction enzyme site and missense variant of p.G191. The HEK293T cell line were screened for template-based repair using AscI digestion and those clones that are positive for the repair were Sanger sequenced for the G191 variant. This process generated three cell lines for these studies: EXOSC3 G191C/+, G191D/G191D, and G191D/+. In brief, HEK293T cells were transfected with plasmids that contain EXOSC3 G191 targeting gRNA, Cas9, and a template DNA containing the G191 missense variant to induce homology directed repair. Additional details are available in ^47^.

### RNA Isolation & Total RNA Sequencing Data Processing

RNA was isolated from four replicate growths of each genotype using the TRIzol reagent-based extraction Zymo Direct-zol RNA prep kit. The RNA was treated with Dnase I on column according to manufacturer’s protocol. The FastQ files from the RNA sequencing were processed based on the following pipeline. First, adapter sequences were trimmed using bbduk tool from BBTools v.38.72 with the following parameters: *k=23 ktrim=r hdist=1 tpe=t tbo=t useshortkmers=t mink=11 qtrim=rl trimq=10 minlength=50*. Reference FASTA file of adapters (*adapter.fa* from the BBTools suite^111^. FastQC v. 0.11.5 was used to determine the quality of trimmed reads^112^. Before indexing, a FASTA reference genome was made by concatenating the Ensembl import GRCh38 version 108 with the ERCC Spike-In sequences. An indexed human reference genome with ERCC Spike-In was made using STAR v.2.7.3a with parameters *-sjdbOverhang 100*^113^. The trimmed reads were aligned to the indexed reference genome using STAR v. 2.7.3a using the parameters *--outFilterType BySJout --outFilterMultimapNmax 20 --alignSJoverhangMin 8 --alignSJDBoverhangMin 1 --outFilterMismatchNmax 999 --outFilterMismatchNoverLmax 0.04 --alignIntronMin 20 --alignIntronMax 1000000 --alignMatesGapMax 1000000 --outSAMattributes NH HI NM MD --outSAMtype BAM SortedByCoordinate*^114^. Gene read counts and junction spanning reads were obtained using the featureCounts tool of Subread v. 2.0.3 with the following parameters accounting for the reverse stranded nature of the library generation: -*p --countReadPairs -s 2* and GTF used in STAR genome indexing previously mentioned^115^.

The gene read count files were imported into RStudio running R v.4.2.3^116,117^. Differential expression analysis was performed using DESeq2 v. 1.44.0^118^. DESeq data set was created using the *DESeqDataSetFromMatrix* function with the design argument *∼ Genotype* (the EXOSC3 genotype) and the reference level to EXOSC3 wildtype genotype. Only genes with total read counts greater than or equal to 5 were included in the estimations of size factors, dispersion, and negative binomial GLM fitting. In the estimation of size factors, the vector of counts aligned to the ERCC Spike-In was used as the *controlGenes* argument in the function *estimateSizeFactors*. A log-fold-change shrink was applied using “*ashr*”^119^. Differential expressed genes were considered to be genes with an FDR-adjusted (Benjamini-Hochberg) *p*-value less than or equal to 0.05. The principal component analysis was conduct using the variance stabilizing transformation of the count matrix and plotting it using the *DESeq2* function *plotPCA*^118^.

The NCBI Gene Expression Omnibus (GEO) was queried to identify RNA-seq datasets involving EXOSC3 knockdown in human cell lines. Two studies were selected: GSE72055 and GSE84172, and the corresponding raw sequencing data (FASTQ files) were retrieved via the SRA. These datasets were processed using the same RNA-seq pipeline described above, with minor modifications made to account for study-specific library preparation chemistries. DESeq2 was used for differential transcript abundance analysis as above.

### RNA-Seq Enrichment Analyses

The list of differential abundant RNA were tested for gene ontology enrichment using the *enrichGO* function of *clusterProfiler* ^120–123^ or Human Protein Atlas enrichment using the *gost* function of *gprofiler2*^58^. The background list was the list of all genes used after the filtering for read counts greater than or equal to 5. The rich factor was calculated where the number of DATs matching to the specific ontology was divided by the number of gene annotated to that ontology. The enrichment data was plotted using a dot plot. To assess the overrepresentation of known UTR elements in differentially abundant transcripts, we obtained manually annotated regulatory sites for trans-factors and cis-elements in UTRs from the Atlas of UTR Regulatory Activity (AURA) version 2.6, filtering for human-specific annotations^124^. Overrepresentation analysis for each regulatory element was performed using Fisher’s exact test, implemented in the R package *stats*, with p-values adjusted using the Benjamini-Hochberg correction. The top 20 most statistically significant regulatory elements were visualized using *ggplot2*^125^.

### Splicing Analysis

To understand the effect of G191 variants on splicing, we utilized *replicate Multivariate Analysis of Transcript Splicing* (rMATS, version 4.1.0) ^71,72,126^. Mapped reads from the total RNA sequencing used the ENSEMBL GRCh38.p13 (hg38) as the annotation for splicing events, including skipped exon (SE), retained intron (RI), mutually exclusive exon (MXE), alternative 5’ splice-site (A5SS), and alternative 3’ splice-site (A3SS). For each splicing event in the annotation, the proportions of inclusion (*I*, the event occurred) and exclusion (*S*, otherwise) were quantified based on the junction reads supporting the respective isoforms in the RNA-seq alignment files of the samples. Changes in the inclusion level between the control and variant samples were estimated to call the differential splicing events, with statistical significance (FDR) evaluated. Alternative splicing events with an FDR < 0.05 in the change of inclusion level were regarded as differentially observed and we focused on splicing phenomenon that suffice the following 1.) having ≥10 support junction reads in all samples of control and mutant, and *2.*) absolute fold-change of inclusion level > 10% between control and variant, were highlighted and forwarded to further analysis. Read counts and junction-spanning reads for EXOSC3 from the RNA sequencing data were visualized using sashimi plots generated with ggsashimi^127^. Junction-spanning reads covering the canonical exons of the first and second isoforms of EXOSC3 (ENST00000327304 and ENST00000396521) were plotted.

### Generating RNA exosome Structural Models

A cryo-EM structure of human RNA exosome (PDB: 6D6Q)^128^ was used to generate a structural model suitable for molecular modelling^75^. Molecular refinement of the RNA exosome protein structure was performed following the Rosetta macromolecular assembly refinement protocol developed by the DiMaio lab^129,130^. To decrease the computational cost of simulating the full RNA exosome complex, a final RNA exosome-bound model was created by truncating all residues outside a 27 Å radius from EXOSC3. The unbound EXOSC3 model was created by extracting EXOSC3 from the full model of the RNA exosome complex. To measure ΔΔ*G*_fold_ with λ-dynamics, an unfolded model of EXOSC3 was made by extracting a pentapeptide centered around G191 (EXOSC3 residues 189-193) from the full RNA exosome model^131^. To create a model of the alternatively spliced EXOSC3 (EXOSC3_alt_), the unbound EXOSC3 model was used and all amino acids after residue 164 were removed. Residues 159-164 were then mutated from VGDLIY to AISSRL using PyMOL’s mutagenesis tool^74^.

All systems, prior to minimization or MD simulations, were solvated in a 0.1 M NaCl solution with the TIP3P water model ^132^ using CHARMM-GUI with a solvent buffer of 20 Å for the unfolded EXOSC3 (pentapeptide) model or 10 Å for all other models. All systems were energy minimized using 1000 to 2000 steps split in stages, starting with restraints on all protein and nucleic acid atoms and then incrementally removing the harmonic restraints such that the final minimization was performed with restraints removed from all atoms, except for the protein backbone atoms located 17 Å away from EXOSC3 and all atoms of the first and last RNA bases. These atoms were restrained to preserve the structure of the RNA exosome complex around EXOSC3 in the truncated model.

### Molecular Dynamics Simulations

MD simulations were performed with the CHARMM molecular software package^133,134^ and the BLaDE interface for running on graphic processing units (GPUs)^135^. The CHARMM36 protein and nucleic acid force fields^136–138^ were used to represent all RNA exosome energetics, and water was represented with the TIP3P water model^132^. All simulations were performed in an isothermal-isobaric ensemble with a pressure of 1.0 atm and a temperature of 30 °C. Long-range electrostatic interactions were modelled with particle mesh Ewald (PME) with force switching and nonbonded cutoffs set to 9 and 10 Å, respectively^76,77,79,139^. The SHAKE algorithm was used to restrain hydrogen atoms^81^. Twelve 300 ns MD simulations were performed of the RNA exosome-bound and unbound native EXOSC3. All other EXOSC3 variants (bound and unbound EXOSC3 G191C, bound and unbound EXOSC3 G191D, and unbound EXOSC3_alt_) were performed for 300 ns in triplicate. The first fifth of each simulation was discarded as equilibration prior to trajectory analysis. For the truncated RNA exosome-bound system, harmonic restraints with a force constant of 10 kcal mol^-1^ Å^-2^ were applied to the backbone atoms of protein residues more than 17 Å away from EXOSC3 and all atoms of the terminal RNA bases. All other atoms remained fully flexible.

### λ-Dynamics Simulations

λD simulations^80,140^ were performed to investigate the thermodynamic changes (ΔΔ*G*_bind_ and ΔΔ*G*_fold_) introduced as a result of the EXOSC3 G191C and G191D mutations. The change in charge accompanying the EXOSC3 G19D perturbation was accounted for using a post-simulation analytic correction (AC)^75^. A routine λD simulation was performed and then a post-alchemical correction^141^ was applied to the ΔΔ*G* results (Eq. 1).

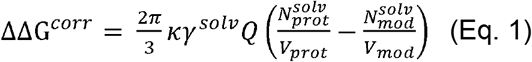

where *k* is the electrostatic constant, *γ^solv^* is the quadrupole moment trace of the solvent model relative to a van der Waals interaction site (calculated as 0.764 e·Å^2^ for the TIP3P water model^78^). Q is the change in charge between alchemical end states. *N^solv^* is the number of solvent molecules, and *V* is the volume of the periodic box averaged over all production frames, where *prot* denotes the RNA exosome-bound EXOSC3 for the binding free energy simulations or the folded EXOSC3 state within the folding free energy simulations and *mod* denotes the unbound EXOSC3 state within the binding free energy simulations or an unfolded EXOSC3 pentapeptide within the folding free energy simulations. Full-residue perturbations were performed, following a strategy developed for alchemical perturbations of proline and glycine residues with λD^142^. The native EXOSC3 G191 was perturbed to Ala, Cys, and Asp residues in a single λD simulation. Ala served as an intermediate state between Gly and Cys or Asp. The Adaptive Landscape Flattening (ALF) algorithm was used to identify appropriate biasing potentials for dynamic sampling of multiple end states^131,143^. Optimal biases were obtained after a cumulative of 200-500 ns of sampling. This large sampling requirement for bias identification likely stems from a combination of the difficulty in sampling both charge-change perturbations and in achieving converged sampling amidst large conformational changes associated with the G191C/D variants. Finally, five replicate λD production simulations were run for the RNA exosome-bound, unbound, and unfolded EXOSC3 systems. The first fifth of each production simulation was excluded as equilibration prior to determination of the free energy changes. A cutoff value of λ ≥ 0.99 was used to classify perturbation end states. Final relative free energy differences were calculated with WHAM^144^, and statistical errors were estimated using bootstrapping.

### Intermolecular Measurements

Structural changes introduced by the EXOSC3 G191C and G191D variants were investigated by analyzing select distances and dihedral angles observed to be disrupted. Three EXOSC3 β_4_-β_L5_ distances of interest (G191 N – Y164 O, G191 O – Y164 N, and I193 N – L162 O) were collected using an inhouse python script. Per-residue average backbone root mean square deviations (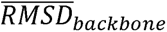) were collected using an inhouse python script following eq. 2:

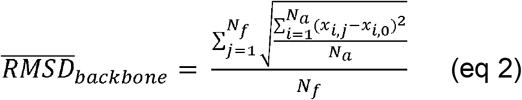

where *N_a_* is the number of backbone atoms in the residue, *N_f_* is the total number of production frames, *x_i,j_* is the coordinates of atom *i* in MD frame *j*, and *x*_*i*,0_ is th coordinates of atom *i* in reference frame 0. The 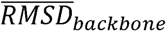 was plotted as the relative change in RMSD with respect to the native G191 (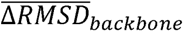) using Maptplotlib^145^.

### Global Proteomics and Proteome Integral Solubility Assays

HEK293T cells with EXOSC3 variants and control were grown in full growth media and subjected to a paired global proteomics and PISA workflow that we have recently reported adapted from prior studies from Gaetani, et. al^146–149^. In brief, Cell pellets were thaweded on ice and resuspended with TPP lysis buffer containing 40 mM HEPES pH 7.5, 200 mM NaCl, 5 mM β-glycerophosphate, 0.1 mM sodium orthovanadate, 2 mM TCEP, 0.4% NP-40, and EDTA-free, cOmplete Protease Inhibitor Cocktail (Roche). The cells were lysed by three rounds of freeze-thaw using liquid nitrogen for freezing and thawing on ice. An aliquot of 100 μg of protein was set aside for global proteomics. The cell lysate was aliquoted into PCR tubes and each aliquot of lysate is heat treated at one of the following temperatures subjected to heat treatment with the following temperatures 25°C, 35°C, 39.3°C, 43.3°C, 48.6°C, 50.1°C, 51.9°C, 54.5°C, 58.7°C, 60.6°C, 74.9°C, and 90°C for 3 minutes followed by rapid cooling to 4°C. After being transferred to microcentrifuge tubes, the heat-treated lysates were centrifugated at 20,000 x g for 30 minutes at 4°C to pellet denatured protein. To prepare samples for Protein Interaction Stability Assay (PISA), 10 μL of the supernatant from each temperature condition was combined into a single tube for replicate of each genotype. For both global proteomics and PISA, proteins were precipitated using trichloroacetic acid (TCA) protocol and digested as previously described ^146,150^. The resulting peptides were desalted with Pierce C18 Spin Columns (ThermoFisher, 89873). Peptides were then labeled with 0.5 mg of tandem mass tag (TMT)pro-16plex label (ThermoFisher) resuspended with anhydrous acetonitrile. The reaction was incubated at room temperature for 1.5 hours. The reaction was quenched with 0.2% hydroxylamine and incubated for 15 minutes at room temperature. Equal volumes of each sample were multiplexed. The resulting peptide mixture was dried via speed vacuuming and desalted with 100 mg C18 SepPak cartridge (Waters) and washed with 5% acetonitrile. The TMTpro-labeled peptides were fractionated with a series of buffers with increasing acetonitrile as described^150^. The fractions were dried down with speed vacuuming and resuspend with 0.1% formic acid.

Nano-LC-MS/MS analyses were performed on an EASY-nLC^TM^ HPLC system (Thermo Fisher Scientific) coupled to Orbitrap Eclipse™ Tribrid™ mass spectrometer (Thermo Fisher Scientific) with a high field asymmetric ion mobility spectrometry (FAIMS) Pro Interface (Thermo Fisher Scientific) at a compensation voltage of -50 V. Each fraction was loaded onto a reversed phase Aurora Ultimate^TM^ C18 ultra-high performance liquid chromatography (UHPLC) column (1.7 μm, 120 Å, 75 μm x 25 cm, IonOpticks) at 400 nL/min. Peptides were eluted from 6-34% with mobile phase B (Mobile phases A: 0.1% formic acid (FA), water; B: 0.1% FA, 80% Acetonitrile (Fisher Scientific) over 110 minutes; 34-80% B for 4 minutes; and dropping from 80-6% B over the final 6 min. The mass spectrometer method was operated in positive ion mode with a 4 sec cycle time data-dependent acquisition method with advanced peak determination and Easy-IC (internal calibrant). Precursor scans (m/z 400-1750) were done with an Orbitrap resolution of 120,000, RF lens% 30, maximum inject time 50 ms, standard automatic gain control (AGC) target, including charges of 2 to 7 for fragmentation with 30 sec dynamic exclusion. MS2 scans were performed with a fixed first mass of 100 m/z, 34% fixed collision energy for HCD fragmentation, 50,000 resolution, 200% normalized AGC target and dynamic maximum IT; the isolation mode was set to quadrupole with an isolation window of 0.7 m/z and without isolation offset.

Resulting RAW files were analyzed in Proteome Discover 2.5 or 3 (Thermo Fisher Scientific) with *Homo sapien* UniProt Reference Proteome FASTA plus common contaminants as previously described^150^. CHIMERYS^151^, MS Amanda 3.0^152^, and SEQUEST-HT^153,154^ searches were conducted with a maximum number of 3 missed cleavages; precursor mass tolerance of 10 ppm; and a fragment mass tolerance of 0.02 Da. Static modifications used for the search were, 1) carbamidomethylation on cysteine (C) residues; 2) TMTpro label on lysine (K) residues and the N-termini of peptides. Dynamic modifications used for the search were oxidation of methionine, phosphorylation on serine, threonine or tyrosine, and protein N-termini dynamic modifications of acetylation, methionine loss, or acetylation with methionine loss on protein N-termini. Percolator False Discovery Rate (FDR) was set to a strict setting of 0.01 and a relaxed setting of 0.05. Values from both unique and razor peptides were used for quantification. In the consensus workflows, peptide TMTpro ion intensities were normalized by total peptide amount with no scaling. Quantification methods utilized lot-specific TMTpro isotopic impurity levels available from Thermo Fisher Scientific. Reporter ion quantification was allowed with S/N threshold of 10 and co-isolation threshold of 30%.

### Site-Directed Mutagenesis for EXOSC3-HaloTag Overexpression Vector

Site-directed mutagenesis (SDM) was performed using the Agilent QuickChange XL Site-Directed Mutagenesis Kit following the manufacturer’s protocol. Mutagenic primers were designed using the Agilent QuickChange Primer Design tool and synthesized. The sequences for the primers are listed in Table 1. The mutagenesis reaction was carried out using a plasmid with an N-terminal HaloTag of EXOSC3 on a pFN21A backbone (Promega, FHC10556) as the template. The PCR reaction was prepared with 50 ng template DNA, 125 ng forward primer, 125 ng reverse primer and other reagent amounts according to manufacturer’s protocol. PCR amplification was performed under the following cycling conditions: Initial denaturation: 95°C for 1 min 30 cycles of amplification with Denaturation: 95°C for 50 s, Annealing: 55°C for 1 min, Extension: 68°C for 6 min, Final extension: 68°C for 7 min. Following PCR, 1 µL of DpnI (10 U/µL) was added to the reaction and incubated at 37°C for 1 hour to digest the parental (non-mutated) plasmid. The digested product was then transformed into XL10-Gold ultracompetent cells (Agilent) by heat shock at 42°C for 45 s, followed by incubation on ice for 2 min. Cells were recovered in 500 µL SOC medium at 37°C for 1 hour with shaking at 225 rpm, then plated on LB-agar plates containing ampicillin (100 µg/mL). After overnight incubation at 37°C, individual colonies were selected for mini-prep plasmid isolation. Mutations were confirmed by Sanger sequencing (Genewiz) using appropriate sequencing primers.

**Table 1:**
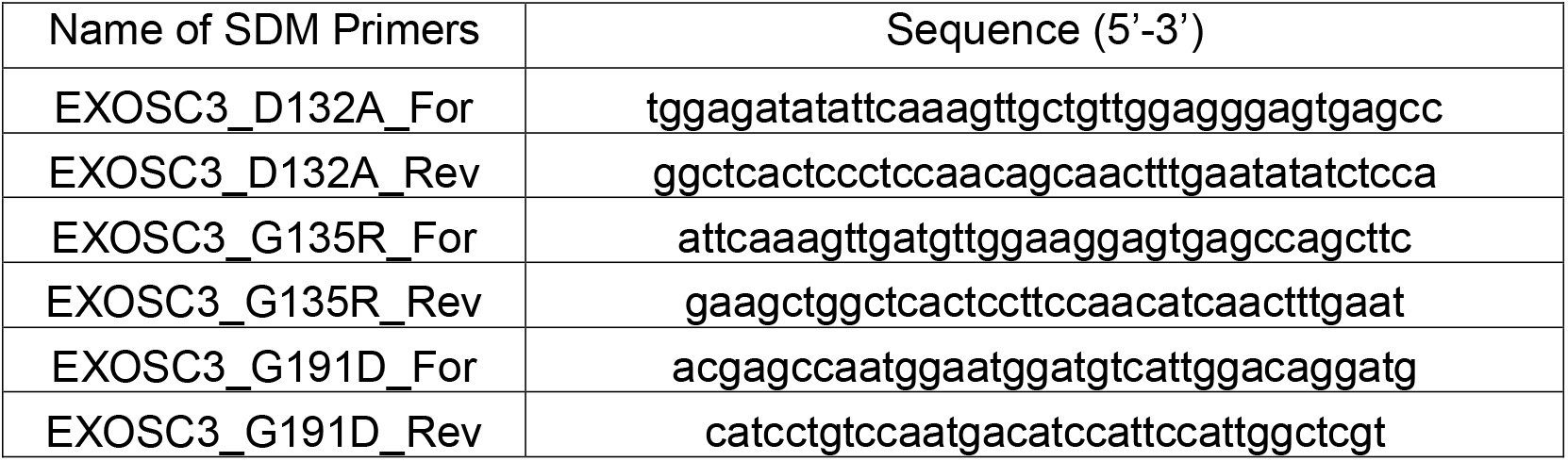
Sequences of primers used for SDM of EXOSC3.

### Transfection

For the EXOSC3-HaloTag overexpression experiments, control and EXOSC3 G191D/G191D cell lines were plated at a density of 3 million cells per T25 flask. The following day, cells were transfected with 4 µg of plasmid DNA encoding either wild-type or G191D variant EXOSC3-HaloTag or a no-DNA control, using polyethylenimine (PEI) transfection reagent. After 72 hours, cells were treated with tetramethylrhodamine (TMR) HaloTag fluorescent ligand and sorted by flow cytometry for positive transfectants.

### Fluorescence-Activated Cell Sorting (FACS) for HaloTag

For HaloTag overexpression experiments, experiments were performed as outlined in ^47^. Briefly, transfected cells were incubated with 5 μM tetramethylrhodamine (TMR) HaloTag ligand (MedChemExpress) in growth media at 37°C for 15 minutes in the dark. Following incubation, cells were washed twice with DPBS and then incubated in fresh growth media for at least 30 minutes before sorting. For HaloTag overexpression experiments (post-TMR ligand labeling), cells were harvested using TrypLE (Thermo Fisher), washed with DPBS (Dulbecco’s phosphate-buffered saline without magnesium or calcium ions), and resuspended in FACS buffer (DPBS supplemented with 0.5% FBS, 1 mM EDTA, and 1 mM HEPES). The cell concentration was adjusted to 7–8 × 10⁶ cells/mL, and samples were passed through a 40 μm cell strainer to remove aggregates. FACS was performed using a Sony MA900 cell sorter equipped with 488 nm and 561 nm laser lines and corresponding emission filters (FL2 585/30 for TMR, FL3 617/30 for mCherry). Data acquisition and sorting were conducted using the Sony MA900 Cell Analyzer software (version 3.1.1). Sorting gates were defined based on forward scatter (FSC), back scatter (BSC), and marker expression (TMR intensity for HaloTag overexpression). Cells were selected based on high FSC-A and BSC-A, with doublets excluded using FSC-H vs. FSC-A gating. After gating, cells with high marker intensities were collected. Target populations were sorted into DPBS using the “Normal” sorting setting. Approximately 700,000 to 1,200,000 marker positive cells were collected. Sorted cells were centrifuged at 1000 × g for 5 minutes at 4°C and resuspended in 8 M urea in 100 mM Tris-HCl (pH 8.5). Samples were either stored at -80°C for future processing or sonicated using the Bioruptor (Diagenode, Denville, NJ) for 30 cycles (30 s on, 30 s off) on high power. Samples were normalized to 50 μg of protein in 100 μL in 8 M Urea in 100 mM Tris-HCl buffer and processed for mass-spectrometry-based global proteomics analysis.

### Pathway Activity Score Analysis

To evaluate the effects of EXOSC3 G191 protein overexpression and EXOSC3 G191D/G191D genotype on protein networks, we quantified pathway activity using TMT-based proteomics data from the overexpression experiment. Protein abundances were normalized using median peptide intensity normalization in ProteomeDiscoverer. Only proteins quantified in all samples with a protein-level FDR ≤ 0.05 were retained for analysis. The resulting abundance matrix, with proteins as rows and samples as columns, was used as input for single-sample gene set enrichment analysis (ssGSEA) to estimate pathway activity scores. EXOSC3 protein abundance values were removed for pathway analysis given experimental design. Gene Ontology (GO) gene sets were retrieved from the MSigDB (Human) using the msigdbr R package^155^, and filtered to include only pathways containing between 10 and 500 quantified proteins. Pathway activity scores were computed using the gsva() function from the GSVA R package^156^, specifying method = "ssgsea". Differential pathway activity across experimental groups (two genotypes by three plasmids with over expression) was assessed using one-way ANOVA followed by post hoc testing. Significant results (ANOVA p ≤ 0.05) were visualized using boxplots generated with the ggplot2 R package ^125,157^.

## Author’s Contributions

- H.R.S.W. - experimental design, data acquisition, data analysis, writing and revision of manuscript
- A.M.R. – experimental design, data acquisition, data analysis, writing and revision of manuscript
- M.P.B. – experimental design, data acquisition, data analysis, writing and revision of manuscript
- R.L. – data analysis
- S.A.P.J. – data analysis, writing and revision of manuscript
- A.H. – data analysis, writing of manuscript
- W.R.S.K. – data acquisition
- J.D.R. – data analysis, data acquisition
- G.S.C. – data acquisition
- S.P. – cell line generation
- E.H.D. – data acquisition
- S. H. L – data analysis, writing of manuscript
- Y.L. – supervision & experimental design
- J.Z.V. – funding acquisition, supervision, experimental design, data analysis, revision of manuscript
- A.L.M. – funding acquisition, project administration, experimental design, data analysis, writing and revision of manuscript

## Acknowledgements

We thank all members of the Mosley lab with special thanks to Neil McCracken, Tara Umberger, and Katelyn Caric for helpful conversations and comments regarding these studies. We also thank Anita Corbett, Homa Ghalei, and Derrick Morton for insightful discussions of this work prior to submission. Sequencing analysis was carried out in the Center for Medical Genomics at Indiana University School of Medicine. The mass spectrometry work was performed by the Center for Proteome Analysis (CPA) and the Mosley lab. Acquisition of the CPA instrumentation used for this project was provided in part by the Indiana University Precision Health Initiative and the IU Simon Comprehensive Cancer Center. The proteomics work was supported, in part, by the Indiana Clinical and Translational Sciences Institute (Award Number UL1TR002529 from NCATS) and, in part, by the IU Simon Comprehensive Cancer Center Support Grant (Award Number P30CA082709 from NCI). We also thank Charles Brandt with the Scalable Compute Archive for work on data sharing and storage solutions for the CPA. This work was supported by the Indiana University Genome Editing Center at Indiana University School of Medicine. This research was supported in part by Lilly Endowment, Inc., through its support for the Indiana University Pervasive Technology Institute. The authors acknowledge the Indiana University Pervasive Technology Institute for providing supercomputing and storage resources that have contributed to the research results reported within this paper. This work was supported in part by Shared University Research grants from IBM, Inc., to Indiana University.

